# Extensive somatic nuclear exchanges shape global populations of the wheat leaf rust pathogen *Puccinia triticina*

**DOI:** 10.1101/2022.11.28.518271

**Authors:** Jana Sperschneider, Tim Hewitt, David C. Lewis, Sambasivam Periyannan, Andrew W. Milgate, Lee T. Hickey, Rohit Mago, Peter N. Dodds, Melania Figueroa

## Abstract

Non-sexual processes such as somatic nuclear exchange are postulated to play a role in the diversity of clonally reproducing dikaryotic rust fungi but have been difficult to detect due to the lack of genome resolution between the two haploid nuclei. We examined three nuclear-phased genome assemblies of *Puccinia triticina*, which causes wheat leaf rust disease. We found that the most recently emerged Australian lineage is derived by nuclear exchange between two pre-existing lineages, which originated in Europe and North America. Haplotype-specific phylogenetic analysis reveal that repeated somatic exchange events have shuffled haploid nuclei between long-term clonal lineages, leading to a global population representing different combinations of a limited number of haploid genomes. Thus, nuclear exchange seems to be the predominant mechanism generating diversity and the emergence of new strains in this otherwise clonal pathogen. Such genomics-accelerated surveillance of pathogen evolution paves the way for more accurate global disease monitoring.

## Introduction

Rust fungi of the order *Pucciniales* are responsible for diseases on some of the most important agricultural crops and threaten food production and ecosystems. For *Puccinia* species causing cereal diseases it is the asexual (uredinial) phase of their life cycles that infects the cereal host, while the sexual phase of the life cycle occurs on different host plants. The population dynamics of these pathogens vary from highly sexual to exclusively clonal depending on the presence and abundance of the alternate host in geographic regions (Figueroa *et al*., 2020). For instance, populations of *Puccinia coronata* f. sp. *avenae* that causes crown rust disease in oats are highly genetically diverse in North America due to the prevalence of the alternate sexual host buckthorn (Nazareno *et al*., 2018; Miller *et al*., 2020). In contrast, *Puccinia graminis* f. sp. *tritici* (*Pgt*) populations that cause stem rust disease in wheat and barley are clonal in most parts of the world due to the scarcity of the alternate host barberry (*Berberis* spp.), but local sexual populations exist in areas where barberry is present (Saunders *et al*., 2019; Patpour *et al*., 2022). Wheat leaf rust disease caused by *Puccinia triticina* (*Pt*) results in substantial crop losses around the world (Kolmer, 2005; Figueroa *et al*., 2017). The sexual host for *Pt, Thalictrum spp*., is scarce in North America and Europe and absent from Australia and is not thought to contribute to population dynamics or diversity of this pathogen (Bolton *et al*., 2008). Molecular genetic analyses of *Pt* populations in different continents indicate that they are composed of relatively few major clonal lineages, with high levels of heterozygosity and linkage disequilibrium and low diversity within lineages consistent with a lack of sexual recombination (Ordoñez & Kolmer, 2009; Ordoñez *et al*., 2010; Kolmer *et al*., 2013; Kolmer, 2019). In Australia, five clonal lineages of *Pt* have been described that appeared to have arisen by exotic incursions to the continent from unknown sources (Huerta-Espino *et al*., 2011; Park, 2015). These include groups defined by the originally detected pathotypes: 53-1,(6),(7),10,11 (detected in 1981); 104-1,2,3,(6),(7),11 (detected in 1984); 76-1,3,5,10,12 (detected in 1996); 10-1,3,9,10,12 (detected in 2004); and 76-3,5,9,10+Lr37 (detected in 2005). These lineages have apparently evolved clonally through stepwise mutations to virulence on single leaf rust resistance genes in wheat to give rise to numerous derived pathotypes (Park, 2015). A further exotic incursion of a lineage with pathotype 104-1,3,4,6,7,9,10,12+Lr37 was detected first in New Zealand in 2012 and subsequently mutated to acquire virulence on *Lr28* (pathotype 104-1,3,4,6,7,9,10,12+Lr37) with the latter pathotype first detected in Australia in 2014 (Warren *et al*., 2018).

In the absence of sexual reproduction, evolution of diversity in rust fungi is limited to mutation and potentially somatic exchange events (Figueroa *et al*., 2020). Numerous laboratory studies have shown that co-infection of a single host plant with two different rust isolates can readily lead to the generation of isolates with novel combinations of virulence genes via somatic exchange (Watson, 1957; Ellingboe, 1961; Flor, 1964; Bartos *et al*., 1969; Lei *et al*., 2017). However, in the absence of high-quality genomic data for each nucleus it was not possible to distinguish between models based on simple exchange of intact haploid nuclei or more complex processes involving genetic recombination between nuclei. Somatic hybridisation can also contribute to rust diversity in the field. For instance, Burdon *et al*. (1981) postulated that a new strain of *Pgt* detected in Australia (where the alternate host is absent) arose via somatic hybridisation based on having a novel combination of virulence and isozyme markers shared with two pre-existing clonal lineages. Similarly, Park *et al*. (1999) proposed somatic hybridisation as the origin of an Australian *Pt* isolate based on shared RAPD (random amplification of polymorphic DNA) and virulence markers. Again, these genetic marker sets were too limited to distinguish between exchange of intact nuclei or recombination. However, recent analysis of fully nuclear-phased genome assemblies clearly demonstrated that somatic exchanges of whole nuclei have contributed to genetic diversity in *Pgt* (Li *et al*., 2019). For instance, one single nucleus-specific haplotype (designated the A haplotype) is nearly identical between *Pgt*21-0 and Ug99, while the second nucleus haplotypes in each isolate (designated B and C in *Pgt*21-0 and Ug99 respectively) are highly divergent. Furthermore, Li et al (2019) also identified three other globally dispersed isolates that contain either the B or C nuclear haplotypes, each with a different second haplotype. Guo *et al*. (2022) used genome admixture analyses to propose that five *Pgt* lineages may be derived by somatic exchange, and identified the putative second parent of Ug99 lineage as another pre-existing African lineage (Clade II).

The detection of nuclear haplotype diversity and somatic hybridisation events requires fully haplotype-phased and nuclear assigned reference genomes, which to date have only been generated for three rust fungi: *P. graminis* f. sp. *tritici* isolate *Pgt*21-0 (Li *et al*., 2019), *P. triticina* isolate *Pt*76 (Duan *et al*., 2022) and *P. coronata* f. sp. *avenae* isolate *Pca*203 (Henningsen *et al*., 2022). Here we generate fully nuclear-phased chromosome assemblies for two additional Australian isolates of *Pt* and use these to compare haplotype diversity across a large set of sequenced *Pt* isolates from around the world. This reveals evidence of extensive nuclear exchange events underlying the origin of major clonal lineages indicating a very substantial contribution of somatic hybridisation to population dynamics.

## Results

### Three haplotype combinations are present in a recent collection of seven Australian *Pt* isolates

Six Australian isolates of *Puccinia triticina* (*Pt*) collected in 2019 and 2020 exhibited four different virulence pathotypes (Table 1). The 19QLD08 isolate shared the same pathotype as the *Pt*76 reference genome isolate (19ACT06) (Duan *et al*., 2022) but with additional virulence for *Lr20*. Both of these are identical to pathotypes found in the lineage derived from pathotype 76-3,5,7,9,10+Lr37, first detected in Australia in 2005 (Huerta-Espino *et al*., 2011; Park, 2015). The 20QLD87 isolate has the same pathotype (104-1,3,4,6,7,8,9,10,12+Lr37) as the lineage first detected in 2014 as an apparent exotic incursion into Australia via New Zealand (Warren *et al*., 2018). The 20ACT90 isolate has the same pathotype as the currently predominant pathotype in Australia, which was first detected in 2016 (Bariana *et al*., 2022), while 19NSW04, 19ACT07 and 20QLD91 share the same pathotype but with additional virulence for *Lr27*/*Lr31* (104-1,3,4,5,6,7,9,10,12+Lr37).

**Table 1:** The *Pt* isolates collected in this study with sample locations in Australia and pathotypes. The nuclear genotypes derived from *k*-mer and lineage assignments according to phylogenetic tree analysis (Figure 1) are shown. The nuclear genotype of 20QLD87 later derived as CD is also shown. (ACT: Australian Capital Territory; QLD: Queensland; NSW: New South Wales).

| Isolate name | Sample collection | Collection year | Pathotype | Nuclear genotype | Lineage |
| --- | --- | --- | --- | --- | --- |
| 19ACT06 (*Pt76*) (Duan *et al.*, 2022) | Canberra, ACT | 2019 | 76-3,5,7,9,10,12,13 | AB | AU1 |
| 19QLD08 | Gatton, QLD | 2019 | 76-1,3,5,7,9,10,12,13 | AB |  |
| 19NSW04 | Wagga Wagga, NSW | 2019 | 104-1,3,4,5,6,7,9,10,12 | BC | AU2 |
| 19ACT07 | Canberra, ACT | 2019 |  | BC |  |
| 20QLD91 | Gatton, QLD | 2020 |  | BC |  |
| 20ACT90 | Canberra, ACT | 2020 | 104-1,3,4,5,7,9,10,12 | BC |  |
| 20QLD87 | Warwick, QLD | 2020 | 104-1,3,4,6,7,8,9,10,12 | CD | AU3 |

The nuclear haplotype similarity of these isolates was assessed by analysis of Illumina genomic sequences against the nuclear-phased *Pt*76 reference chromosome assembly (Duan *et al*., 2022). A *k-*mer containment analysis (Ondov *et al*., 2019) showed that the *Pt*76 A and B nuclear haplotypes are both fully contained (100%) in the Illumina reads from this isolate itself as well as those of the 19QLD08 isolate (Figure 1A), which confirms that these are members of the same clonal lineage. However, while the *Pt*76 B haplotype is fully contained in the Illumina reads of the 20ACT90, 20QLD91, 19ACT07 and 19NSW04 isolates, the A haplotype is not, suggesting that these haplotypes may share the B nuclear haplotype, in combination with another divergent haplotype (hereafter named C). Neither the *Pt*76 A nor B haplotypes are fully contained in reads of 20QLD87, suggesting a different unknown genomic composition. To further confirm this, we aligned the Illumina reads against the *Pt*76 reference and generated maximum likelihood phylogenetic trees based on SNPs called against the combined A and B haplotypes or the individual A and B haplotypes separately. The combined A/B haplotype phylogenetic tree showed that these isolates represent three distinct lineages (Figure 1B); AU1 comprising 19ACT06 and 19QLD08 (AB haplotype), AU2 comprising 20ACT90, 19NSW04, 19ACT07 and 20QLD91 (all BC haplotype), and 20QLD87 being a singleton lineage designated AU3 (unknown haplotypes). The AU1 isolates remained separated in a phylogenetic tree derived from the A haploid genome, confirming that his haplotype is not shared with the other isolates (Figure 1C). However, the AU1 and AU2 isolates grouped together in single closely related clade in a phylogenetic tree based on only the B genome SNPs, indicating that they share the B haplotype, consistent with the *k*-mer containment analysis (Figure 1D). Thus, at least three haplotype combinations (AB, BC and a third unknown) are present in this recent collection of Australian *Pt* isolates.

**Figure 1:**
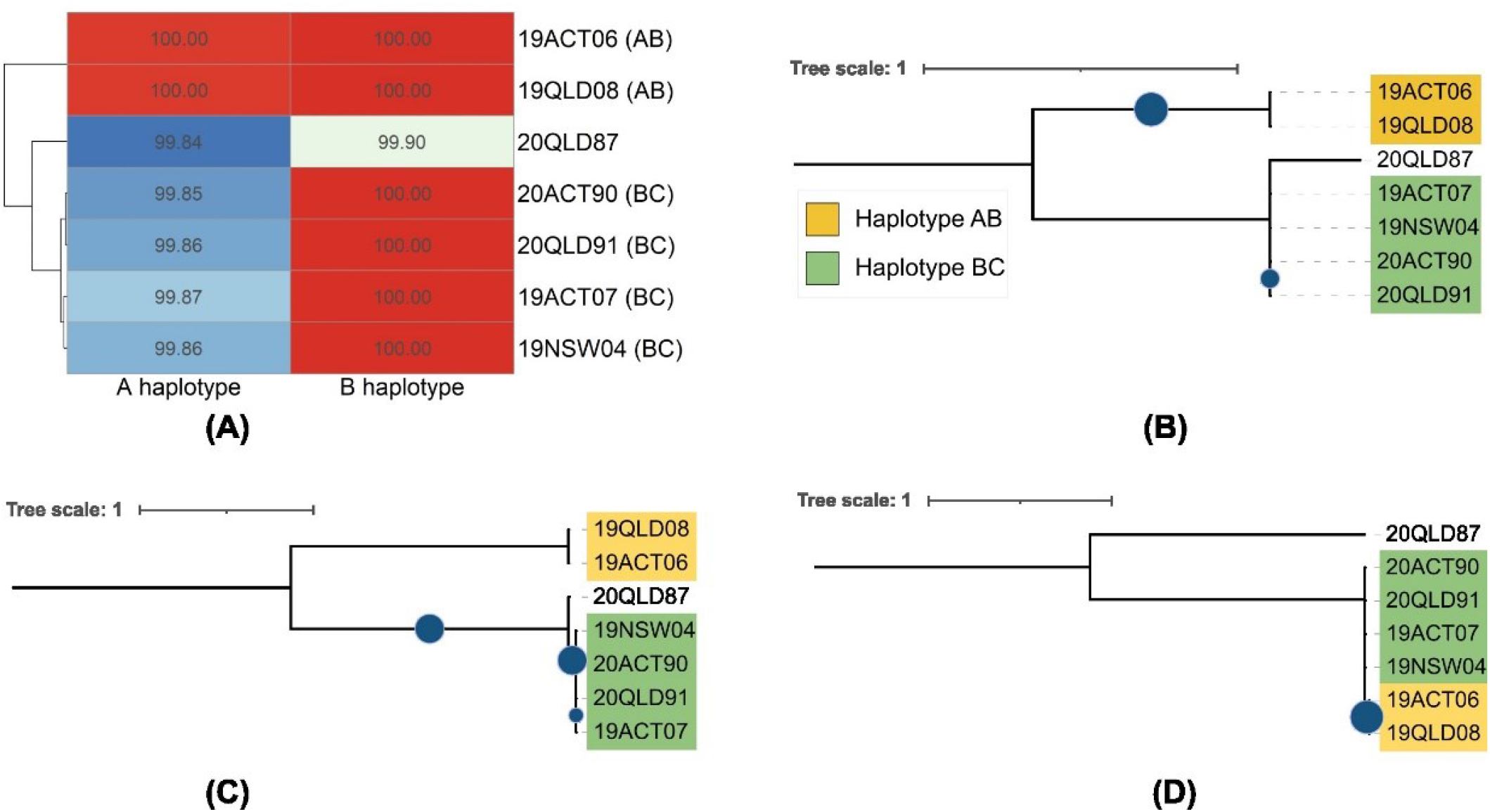
Three distinct lineages and four haplotypes are present in a collection of seven Australian *Pt* isolates. **(A)** *k*-mer genome containment scores against the *Pt*76 (19ACT06) haplotypes cluster into three haplotype groups. A value of 100 indicates that the haplotype genome is fully contained in the sequencing reads of an isolate. **(B)** The phylogenetic tree against the combined haplotypes *Pt*76 A and B indicate three lineages. **(C)** The phylogenetic tree against the *Pt*76 haplotype A shows that two isolates share the A haplotype. **(D)** The phylogenetic tree against the *Pt*76 haplotype B shows that six isolates share the B haplotype. Bootstrap values of over 80% are indicated with blue circles.

### Hi-C integration during HiFi genome assembly returns phased haplotypes in dikaryons with minimal phase switching

To further analyse haplotype similarity in these isolates, we generated nuclear-phased genome assemblies for 19NSW04 (presumed BC haplotype) and 20QLD87 (unknown haplotype) as representatives of the two additional detected lineages. Hifiasm (Cheng *et al*., 2021) assembly of PacBio HiFi long-read sequences and incorporating Hi-C sequencing data returned genome assemblies for each isolate separated into two haplotypes. Each raw haplotype assembly was 123-129 Mb in size, highly contiguous (L50 > 6 Mb) and with BUSCO completeness of over 95% (< 5% duplicated) (Supplementary Table S1). For diploid organisms, hifiasm with Hi-C integration can phase entire chromosomes, but the two haplotype assemblies contain a mix of parental chromosomes as both haplotypes are present in one cell. In contrast, in dikaryons the haplotypes are physically separated in individual nuclei and it was not clear whether the nuclear signal in the Hi-C would affect the hifiasm haplotype assignment. We therefore used the NuclearPhaser pipeline (Duan *et al*., 2022) to assess nuclear phasing in these assemblies using the Hi-C data. In both cases the haplotype1 and haplotype2 assemblies are close to perfectly nuclear-assigned, with only two contigs larger than 150 Kb (1.2 Mb in total) assigned to the incorrect phase (Supplementary Figure S1) in 19NSW04 and only a single mis-assigned contig (2.2 Mb) in 20QLD87 (Supplementary Figure S1). After re-assigning these contigs, ∼92% and ∼85% of *trans* Hi-C contacts occur within a haplotype in 19NSW04 and 20QLD87, respectively. This indicates that the majority of contigs in the two assemblies are fully-phased but minor phase switches within contigs could still be present, particularly for 20QLD87. NuclearPhaser did not detect any phase switches in contigs in the 19NSW04 assembly, while potential phase switches were detected in three contigs in the 20QLD87 assembly. In previous Canu or HiCanu-based assemblies (Koren *et al*., 2017; Nurk *et al*., 2020) we observed that phase switches occurred at haplotig boundaries (Li *et al*., 2019; Henningsen *et al*., 2022) as these assemblers tend to break contigs at points of phase ambiguity. However, this was not the case in these hifiasm contigs, which are generated by a different assembly process (Cheng *et al*., 2021). We therefore assembled the 20QLD87 HiFi reads with HiCanu and found that the predicted phase switch regions in all three contigs corresponded to HiCanu haplotig boundaries, and the HiCanu haplotigs at that boundary also switch phase (Figure 2). Thus, we selected the HiCanu alignment coordinates as breakpoints to correct these phase switches.

**Figure 2:**
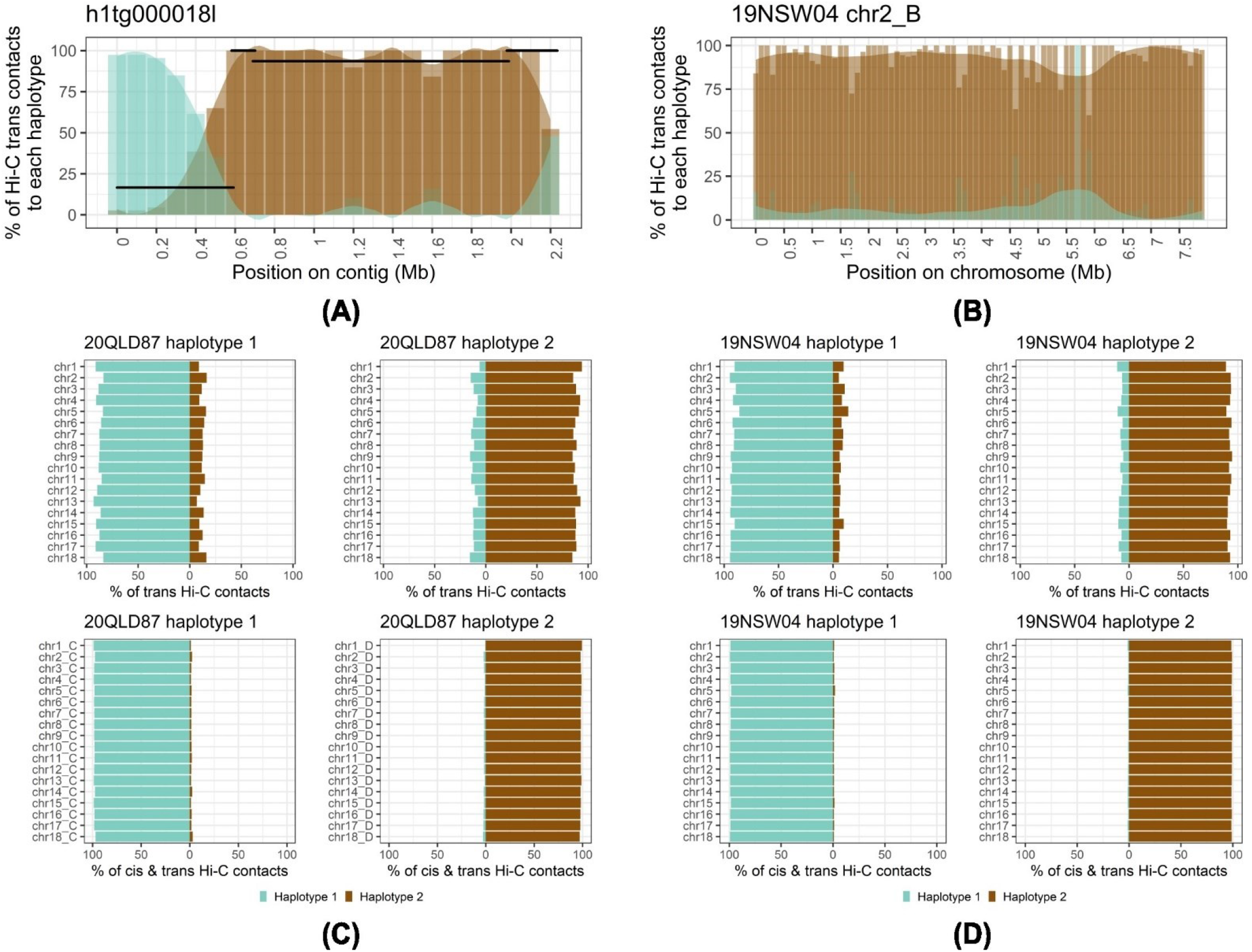
Percentage of Hi-C contacts that link within and between nuclear-separated haplotypes. **(A)** The % of Hi-C *trans* contacts that link to haplotype 1 (turquoise) and 2 (brown) in each 100kbp bin (with an associated smoothing lines) are shown for hifiasm contig h1tg000018l, which contains a phase switch. HiCanu haplotig alignment positions (black segments) are shown at the y-coordinate that corresponds to their Hi-C contacts to haplotype 1. **(B)** The % of Hi-C *trans* contacts that link to each haplotype for the fully phased 19NSW04 chromosome 2B. **(C and D)** Graphs showing the % of Hi-C *trans* contacts (upper panel) or *cis* and *trans* contacts (lower panel) that link to haplotype 1 (turquoise) and 2 (brown) for the 20QLD87 (C) and 19NSW04 (D) chromosome assemblies.

Contig scaffolding resulted in 18 chromosomes assigned to each nuclear haplotype in both 19NSW04 and 20QLD87 assemblies, with over 90% of *trans* Hi-C links occurring within a nucleus as expected for dikaryons (Figure 2, Supplementary Figure S2). The four chromosome haplotype assemblies range from 121.8 Mb to 123.3 Mb in length (Table 2), similar to those of the previously sequenced *Pt76* isolate (121.6 Mb and 123.9 Mb) (Duan *et al*., 2022). Additional unplaced contigs from 19NSW04 and 20QLD87 are mainly small (L50 of 76.2 Kb and 38.7 Kb respectively) with few genes and are highly enriched for repetitive sequences, especially ribosomal RNAs (Table 2). We annotated genes in 19NSW04 and 20QLD87 with a pipeline optimised for effector annotation and re-annotated *Pt*76 with the same pipeline. This increased the number of genes in *Pt*76 from a previously reported 29,052 (Duan *et al*., 2022) to 36,343 (haplotype A: 17,958 genes; haplotype B: 17,813 genes; unplaced contigs: 572 genes), which is in line with the predicted gene numbers in 19NSW04 and 20QLD87 (36,322 and 37,846, respectively; Table 2), as well as with reported gene numbers for *P. graminis* f. sp. *tritici* (*Pgt*21-0: 36,311 genes (Li *et al*., 2019)).

**Table 2:**
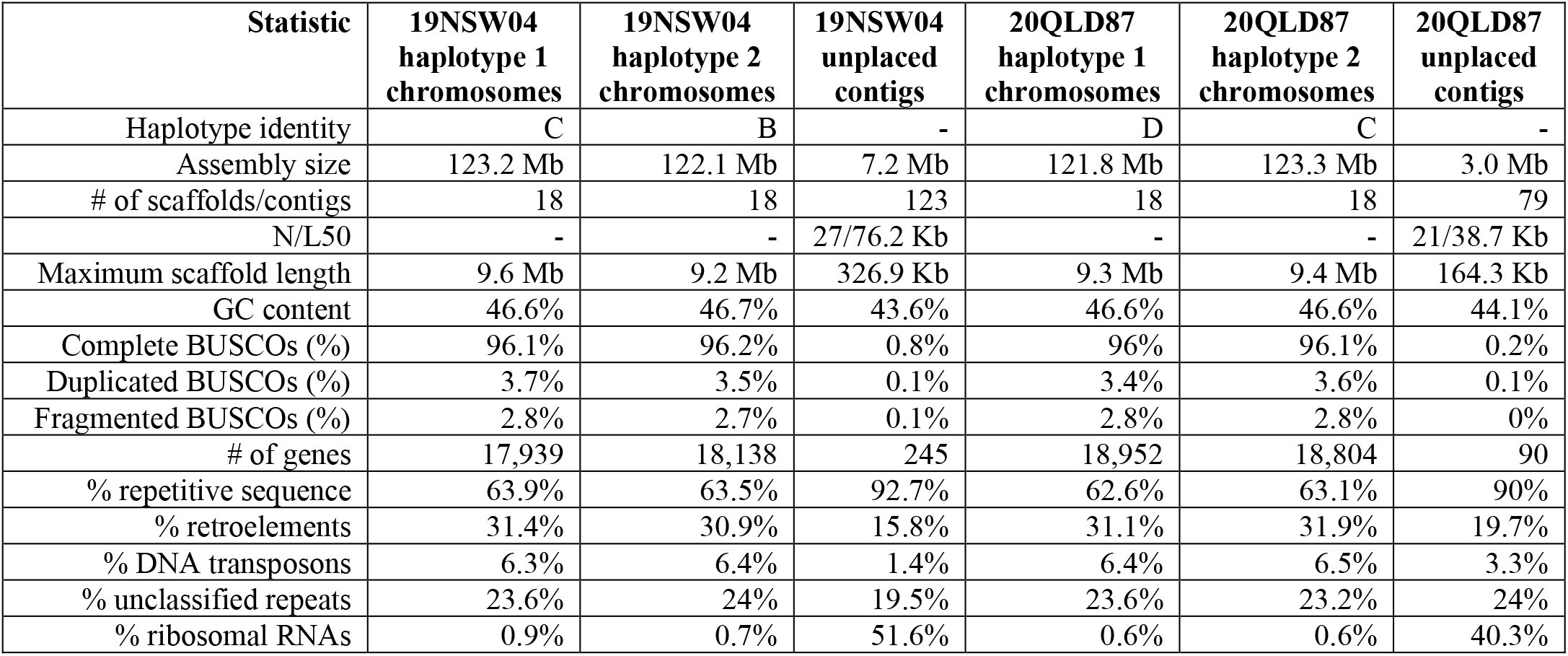
Assembly statistics for the two haplotypes of the scaffolded hifiasm assemblies with Hi-C integration.

| Statistic | 19NSW04<br>haplotype 1<br>chromosomes | 19NSW04<br>haplotype 2<br>chromosomes | 19NSW04<br>unplaced<br>contigs | 20QLD87<br>haplotype 1<br>chromosomes | 20QLD87<br>haplotype 2<br>chromosomes | 20QLD87<br>unplaced<br>contigs |
| --- | --- | --- | --- | --- | --- | --- |
| Haplotype identity | C | B | - | D | C | - |
| Assembly size | 123.2 Mb | 122.1 Mb | 7.2 Mb | 121.8 Mb | 123.3 Mb | 3.0 Mb |
| # of scaffolds/contigs | 18 | 18 | 123 | 18 | 18 | 79 |
| N/L50 | - | - | 27/76.2 Kb | - | - | 21/38.7 Kb |
| Maximum scaffold length | 9.6 Mb | 9.2 Mb | 326.9 Kb | 9.3 Mb | 9.4 Mb | 164.3 Kb |
| GC content | 46.6% | 46.7% | 43.6% | 46.6% | 46.6% | 44.1% |
| Complete BUSCOs (%) | 96.1% | 96.2% | 0.8% | 96% | 96.1% | 0.2% |
| Duplicated BUSCOs (%) | 3.7% | 3.5% | 0.1% | 3.4% | 3.6% | 0.1% |
| Fragmented BUSCOs (%) | 2.8% | 2.7% | 0.1% | 2.8% | 2.8% | 0% |
| # of genes | 17,939 | 18,138 | 245 | 18,952 | 18,804 | 90 |
| % repetitive sequence | 63.9% | 63.5% | 92.7% | 62.6% | 63.1% | 90% |
| % retroelements | 31.4% | 30.9% | 15.8% | 31.1% | 31.9% | 19.7% |
| % DNA transposons | 6.3% | 6.4% | 1.4% | 6.4% | 6.5% | 3.3% |
| % unclassified repeats | 23.6% | 24% | 19.5% | 23.6% | 23.2% | 24% |
| % ribosomal RNAs | 0.9% | 0.7% | 51.6% | 0.6% | 0.6% | 40.3% |

### *Pt* isolate 19NSW04 shares one nuclear haplotype with each of the isolates *Pt*76 and 20QLD87

Genome sequence alignment of the fully-phased haplotypes of 19NSW04, 20QLD87 and *Pt*76 showed that within each isolate, the two separate haplotypes have average sequence identity of 99.50% (divergence 0.50%), with ∼303,000 to 334,000 distinguishing SNPs (Figure 3A, Supplementary Table S2). Similarly, the *Pt*76 A haplotype and 20QLD87 haplotype1 showed average sequence identity of 99.49% with ∼328,000 SNPs. However, the 19NSW04 haplotype2 chromosomes share remarkably high sequence similarity with the *Pt76* B chromosomes with only 2,966 SNPs and average sequence alignment identity of 99.99% (divergence 0.01%). In addition, the 19NSW04 haplotype1 chromosomes share similarly high sequence identity (99.99%) with the 20QLD87 haplotype2 chromosomes, with only 2,226 SNPs. Thus, we assigned the 19NSW04 haplotype2 as B, the 19NSW04 haplotype1 and 20QLD87 haplotype2 as C and the 20QLD87 haplotype1 as D (Table 2, Figure 3A). The 19NSW04 haplotype C assembly contains a translocation between chromosomes 2 and 6, which is not present in any of the other haplotypes, including the 20QLD87 C haplotype (Figure 3B). Visual inspection of HiFi read coverage at the translocation breakpoints and the Hi-C contact maps (Supplementary Figure S2) confirmed that the translocation does not result from mis-assembly (data not shown). Thus, *Pt76* and 19NSW04 share a near-identical copy of haplotype B and 19NSW04 and 20QLD87 share a near-identical copy of haplotype C, with a single translocation in the 19NSW04 C haplotype. These data suggest that nuclear exchange underlies the relationship between these isolates, with the simplest scenario being that the BC lineage arose by a somatic hybridisation event between isolates of the AB and CD lineages.

**Figure 3:**
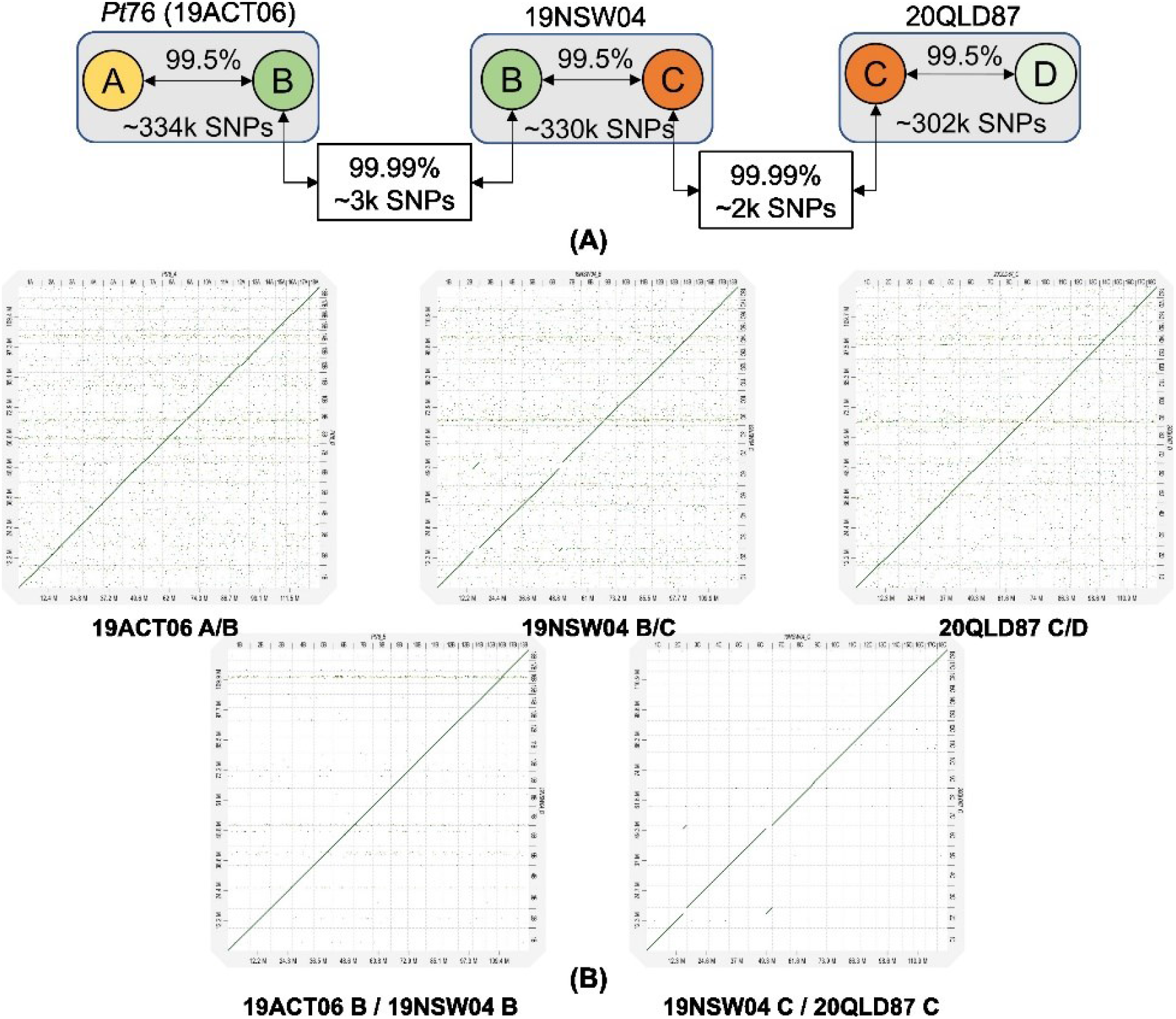
*Pt76* (19ACT06) and 19NSW04 share a near-identical copy of haplotype B and 19NSW04 and 20QLD87 share a near-identical copy of haplotype C. **(A)** Diagram showing the average identity of genomic alignments and total number of SNPs among haplotypes of 19ACT06, 19NSW04 and 20QLD87. **(B)** Dot plot genomic alignments show a single translocation in the 19NSW04 C haplotype.

### Clonal lineages with the AB and CD nuclear haplotypes occur in Europe and North America, respectively

To assess haplotype relationships amongst global *Pt* isolates, we used whole-genome sequencing data from three additional data sources including 20 isolates from Australia and New Zealand (Wu *et al*., 2017), seven isolates from Australia (Wu *et al*., 2020) and 120 worldwide isolates mostly from North America and Europe (Fellers *et al*., 2021) (Supplementary Data S1). Sequencing reads of these 154 isolates were aligned against the three phased diploid genomes and SNPs called against the combined diploid genomes or the individual haploid haplotypes were used to construct phylogenetic trees. The phylogeny derived from SNPs against the diploid 19NSW04 reference genome (Figure 4) showed a very similar topology to a previously reported phylogeny for the 120 global isolates and largely confirms the placement of the North American isolates into six clades (Fellers *et al*., 2021). However, five isolates originally classified in the North American groups NA1 (99NC), NA2 (04GA88-03), NA3 (11US116-1 and 11US019-2) and NA5 (84MN526-2) did not cluster with the rest of these groups in this phylogenetic tree or subsequent trees (see below). Inspection of the phylogeny in Fellers et *al*. (2021) revealed that these isolates were basal to and significantly diverged from these clusters, consistent with these representing different lineages. We classified the 11US116-1 and 11US019-2 isolates as a separate clade NA7 based on this data and results below which indicate that they contain a novel haplotype combination relevant to the evolution of the North American population.

**Figure 4:**
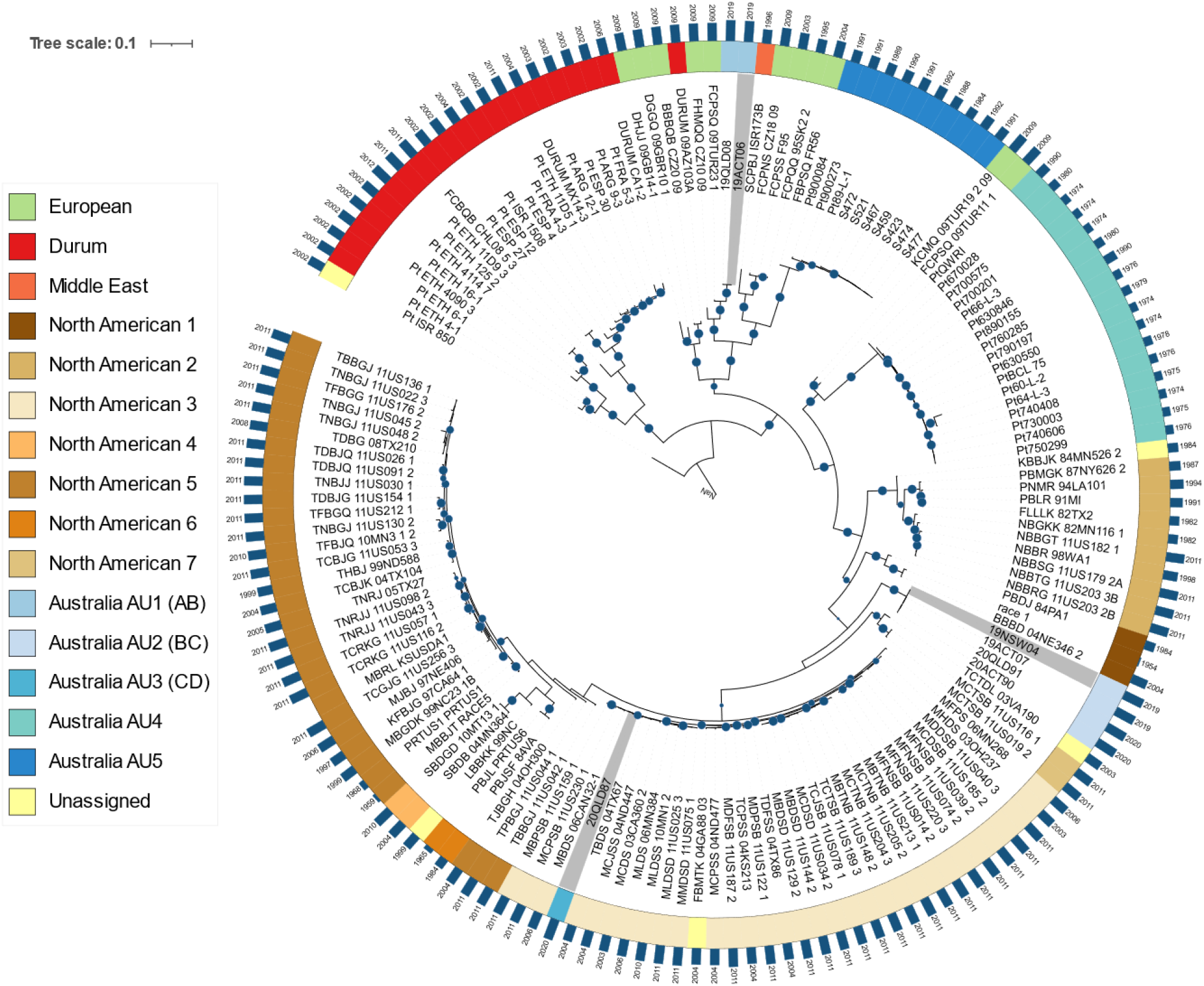
Phylogenetic tree with the diploid genome of 19NSW04 (BC) as the reference. A maximum likelihood tree was generated from 175,974 bi-allelic SNPs. Isolates from the five Australian *Pt* clades (AU1 to AU5), seven North American clades (NA1-NA7) or Europe are indicated by colour according to the legend inset. Five isolates are unassigned to clades, i.e. Pt_ISR_850 (*Aegilops speltoides*, outgroup), 03VA190 (unassigned in Fellers *et al*. (2021) and three others which are in disagreement with the clade assignment given in Fellers *et al*. (2021). *Pt* isolates with fully phased haplotype genome references are highlighted in grey. Bootstrap values over 80% are indicated with blue circles. The year of collection for each isolate is shown next to the blue bars.

The older Australian isolates cluster in two clonal groups separate from the recent isolates (Figure 4): one designated AU4 containing isolates collected between 1974 and 1990; and another designated AU5 containing isolates collected between 1984 and 1992 and representing the clonal lineage derived 104-1,2,3,(6),(7),11 first detected as an exotic incursion in 1984 (Wu *et al*., 2017). The final Australian isolate, 20QLD87 (AU3; CD haplotype) was placed within the NA3 clonal group of North American isolates in this phylogenetic tree indicating that it represents a clonal lineage that arrived in Australia as a result of intercontinental migration. Also of note is that the AU5 group is closely related to a French isolate (FR56) collected in 2004 and the AU1 (AB) lineage closely groups with a Turkish isolate collected in 2009 (09TUR23-1). These two isolates had previously been placed in the European groups EU7 and EU2, respectively, (Kolmer *et al*., 2013), indicating that these two lineages are common to both Australia and Europe.

### Clonal lineages of *Pt* share common nuclear haplotypes in different combinations

We also constructed phylogenies using SNPs from the individual A, B, C and D haplotypes to identify lineages sharing these haplotypes (Figure 5, Supplementary Figures S3-S6). In a phylogenetic tree derived from the A haplotype SNPs (Figure 5A, Supplementary Figure S3), the AU1 isolates (AB) again form a clonal clade with the Turkish isolate (09TUR23-1, EU2), but also with an isolate collected in 2009 from Czech-Slovakia (CZ10-09, EU5), as well as with the AU5 group and the closely related FR56 isolate (EU7 group), suggesting that these groups all share a nucleus with very high similarity to the A haplotype of *Pt*76. In a phylogeny derived from the B haplotype (Figure 5B, Supplementary Figure S4) the AU1 (AB) and AU2 (BC) groups form a clonal clade with isolate 09TUR23-1, again confirming that this EU2 isolate contains both the A and B haplotypes.

**Figure 5:**
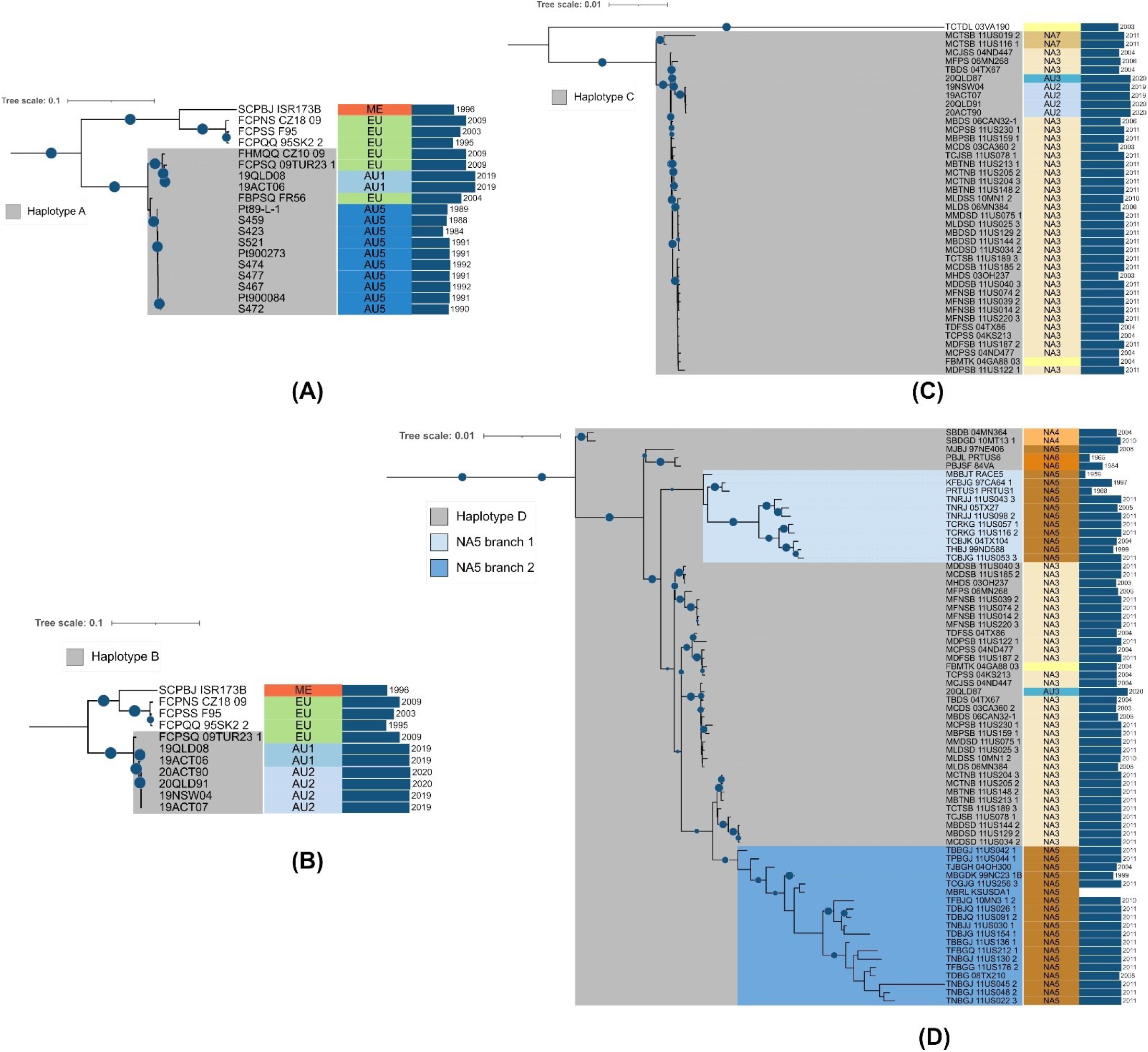
Pruned phylogenetic trees of global *Pt* isolates against the individual A, B, C and D haplotypes. Phylogenetic trees were constructed based on SNPs called against the single haplotypes (Supplementary Figures S3 to S6) and sub-branches of the trees containing isolates with the relevant haplotype reference are displayed. (A) *Pt76* haplotype A; (B) 19NSW04 haplotype B; (C) 20QLD87 haplotype C; and (D) 20QLD87 haplotype D. Bootstrap values over 80% are indicated with blue circles. Clades are indicated next to the name of each isolate (ME: Middle East, EU: European, AU1-AU5: Australian 1-5, NA1-7: North American 1-7). The year of collection for each isolate is shown next to the blue bars.

In the C haplotype derived phylogenetic tree (Figure 5C, Supplementary Figure S5), the AU2 (BC) and 20QLD87 (CD) isolates form a clonal group with isolates from the North American clade 3 (NA3), which again confirms that the C haplotype is shared between these lineages. The C haplotypes of the AU2 isolates are most closely related to 20QLD87, which is consistent with 20QLD87 representing the parental lineage that donated the C nucleus to this hybrid lineage. Likewise, 20QLD87 (CD) again formed a clonal group including the NA3 isolates in a phylogenetic tree derived from SNPs detected against the D haplotype (Figure 5D, Supplementary Figure S6). However, this group also included isolates from the North American clades NA4, NA5 and NA6, suggesting that they all share a common D haplotype. The NA3 group (CD) branches from within the NA5 group, indicating that the D genome in the NA3 lineage is likely derived from a parental isolate from NA5. The NA4 and NA6 groups branch from older nodes in this clade, indicating that their D genomes diverged earlier. In addition, three other North American isolates (99NC, 03VA190, 84MN526_2) that form singleton branches in the other phylogenetic trees were closely related and basal to this D genome-containing group, suggesting they may contain versions of the D haplotype with even older divergence times (Supplementary Figure S6). Notably, the two isolates that we designated as group NA7 (11US116-1 and 11US019-2) cluster with the 20QLD87 and NA3 isolates in the C haplotype derived phylogenetic tree, but not in the D haplotype or other phylogenies suggesting that they share the C haplotype only (Figure 5C-D). These two isolates are basal to the NA3 clade in the C haplotype tree with strong bootstrap support for this node, consistent with these isolates representing the other parental lineage of the NA3 (CD) hybrid. Thus, the data support that the NA3 lineage is derived by a nuclear exchange event between isolates of NA7 and NA5, which donated the C and D nuclear haplotypes, respectively. However, close examination of the D genome phylogenetic tree indicates that the NA-5 group is divided into two separate branches with strong bootstrap support (Figure 5D). Branch 1 is older and ancestral to NA3 consistent with being the D haplotype donor. On the other hand, branch 2 diverged more recently from within the NA3 group. This suggests that a subsequent nuclear exchange event may have occurred in which the D genome of an NA3 isolate was swapped back into an NA5 isolate to recreate a similar haplotype combination but with a different evolutionary history.

A *k-*mer containment analysis (Table 3, Supplementary Data S1) confirmed the haplotype genome contents and relationships determined from the phylogenies above. Sequencing reads of isolate 09TUR23-1 fully contain the *k*-mer sets of both the A and B nuclear haplotype references, but not those of the C or D haplotypes. Isolate CZ10-09 fully contains the A haplotype, while FR56 and the AU5 group are both very closely related to the A haplotype with 99.99% *k*-mer containment. The NA3 isolates fully contain the C and D haplotypes, while NA7 isolates contain complete representations of the C haplotype only. Likewise, the NA6 isolates fully contain the D haplotype and the NA4 and NA5 are both very closely related to the D haplotype with 99.99% *k*-mer containment. Taken together, numerous clonal lineages share these four nuclear haplotypes in various combinations, suggesting that somatic nuclear exchange is a common occurrence in natural *Pt* populations.

**Table 3:** *k*-mer genome containment scores against sequencing reads of various *Pt* isolates and clades.

| *Pt* isolate/clades | *k*-mer genome containment score |  |  |  | Haplotypes present |
| --- | --- | --- | --- | --- | --- |
|  | *Pt*76<br>haplotype A | 19NSW04<br>haplotype B | 19NSW04<br>haplotype C | 20QLD87<br>haplotype D |  |
| AU1 | 100% | 100% | 99.78% | 99.91% | AB |
| AU2 | 99.86% | 100% | 100% | 99.97% | BC |
| 20QLD87 (AU3) | 99.84% | 99.90% | 100% | 100% | CD |
| AU5 | 99.99% | 99.90% | 99.85% | 99.91% | A |
| AU4 | 99.88% | 99.90% | 99.85% | 99.92% | Unidentified |
| 09TUR23 1 | 100% | 100% | 99.88% | 99.97% | AB |
| FR56 | 99.99% | 99.96% | 99.92% | 99.97% | A |
| CZ10 09 | 100% | 99.95% | 99.92% | 99.98% | A |
| NA1 | 99.88% | 99.93% | 99.90% | 99.97% | Unidentified |
| NA2 | 99.90% | 99.92% | 99.91% | 99.94% | Unidentified |
| NA3 | 99.83% | 99.89% | **100%** | **100%** | CD |
| NA4 | 99.76% | 99.87% | 99.73% | **99.99%** | D |
| NA5 branch 1 | 99.83% | 99.89% | 99.88% | **99.99%** | D |
| NA5 branch 2 | 99.85% | 99.89% | 99.91% | **99.99%** | D |
| NA6 | 99.83% | 99.91% | 99.85% | **100%** | D |
| NA7 | 99.84% | 99.89% | **100%** | 99.94% | C |
| Durum | 99.76% | 99.65% | 99.72% | 99.67% | Unidentified |
| Middle East | 99.84% | 99.92% | 99.75% | 99.90% | Unidentified |
| Other European isolates | 99.91% | 99.95% | 99.85% | 99.94% | Unidentified |

Because the whole genome sequence data used above is biased towards North American and Australian isolates, we combined this with a data set derived from restriction site-associated genotyping by sequencing (GBS) SNP analysis of 559 isolates representing 11 global regions (North America, South America, Middle East, Central Asia, Europe, East Africa, Russia, China, Pakistan, New Zealand, and South Africa) (Kolmer *et al*., 2020). Since the GBS SNP data was derived from mapping to the *Pt* ASM15152v1 draft assembly (Kolmer *et al*., 2020), we mapped the whole genome sequence data of the 154 isolates onto this reference and extracted SNP genotypes on a set of 631 polymorphic sites that are represented in both data sets. A phylogenetic tree constructed from this data (Supplementary Figure S7) showed an overall similar topology to the whole genome tree (Figure 4) for the *Pt* isolates and regional clades common to both data sets, confirming that this analysis with a reduced SNP set is robust.

This analysis confirmed that the AU1 isolates (AB haplotype) group into a single clade containing not only the 09TUR23-1 isolate, but also all of the other isolates (19) belonging to the EU2 lineage, confirming the shared genotype of AU1 and EU2 (Figure 6A). In addition, isolates of the Central Asian clade CA1 and Pakistan clade PK3 fall within this lineage group, suggesting that this lineage is common to Europe, Asia and Australasia. The 20QLD87 (CD haplotype) was again placed with the NA3 clonal group as well as with isolates from the European EU8 (4 isolates) and South American SA3 clades (22 isolates) (Figure 6B). This indicates that this clonal lineage is common to the Americas, Europe and Australasia. However, the previously defined EU8 clade is split into two groups in this tree, with the second group (11 isolates) forming a clonal group with the AU2 (BC) isolates and some isolates from Pakistan (PK-2 clade). The shared C genome between these two EU8 subgroups may explain why they were not separated based on SSR analysis (Kolmer et al 2013). Importantly, this suggests that there may have been hybrid BC haplotype isolates in Europe in 2009 before they were detected in Australia, indicating either independent hybridisation events in both continents or migration of a hybrid strain from Europe to Australia. This phylogenetic tree also supported the clonal relationship between the AU5 group and all eight isolates of the EU7 clade represented by FR56 in the whole genome phylogenetic trees, consistent with the introduction of this lineage to Australia from Europe (Figure 6C).

**Figure 6:**
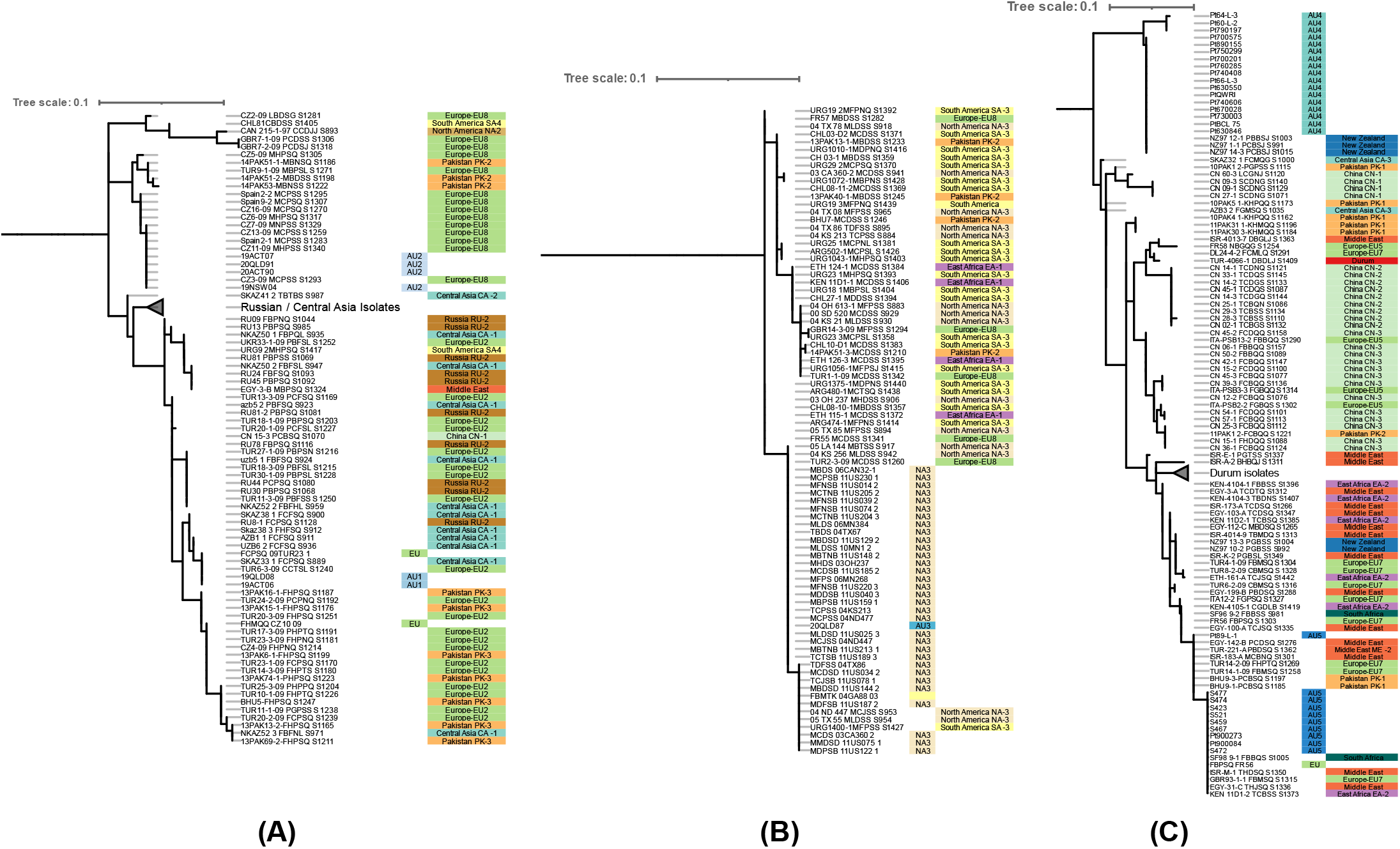
Pruned phylogenetic trees of SNP genotypes shared between GBS and whole-genome data. **(A)** The part of the tree that contains the Australian AU1 isolates also contains the EU2 lineage as well as clades from Central Asia and Pakistan, suggesting this lineage is common to this broad geographical region. The Australian AU2 isolates cluster with one group of the European clade EU8. **(B)** The Australian AU3 isolate is again placed with the North American clade NA3. **(C)** The Australian AU5 isolates group with the EU7 lineage, whereas the AU4 isolates group only with three isolates from New Zealand. Full tree is shown in Supplementary Figure S7.

### Genetic diversity of the mating type loci in *P. triticina*

Mating compatibility in many basidiomycetes is controlled by two loci. The *a* locus encodes a pheromone/receptor pair and the *b* locus encodes two divergently transcribed homeodomain transcription factors known as bEast and bWest (bE and bW) (Bakkeren & Kronstad, 1993). Two alleles of the *a* and *b* locus genes were previously described in the Race 1 (BBBD) reference genome of *Pt* (Cuomo *et al*., 2017). The two alleles (+ and -) of the *a* locus receptor (STE3.2 and STE3.3 genes respectively) are both present and identical in each of our assembled genome references on chromosome 4, with the B and D haplotypes encoding the + allele and the A and C haplotypes the - allele. Examination of SNP calling data against the *Pt*76 diploid reference genome revealed that the 154 *Pt* isolates with genome sequence data all contained both genes with no more than one or two SNPs (Supplementary Data S1). Given that STE3.2 and STE3.3 only share 49% identity, these data confirm that there are only two allelic variants of this locus in this population with all isolates heterozygous. Given that many of these isolates are related by hybridisation and nuclear exchange events, this is consistent with successful dikaryon formation after somatic hybridisation requiring the presence of different *a* locus alleles in the two nuclear haplotypes, as is expected for sexual compatibility of gametes.

The *b* locus encoding the bE and bW proteins is located on chromosome 9, but in contrast to the *a* locus, multiple divergent alleles were detected, with the A haplotype containing the same *b2* allele defined from *de novo* RNAseq assemblies in race 1 (Cuomo *et al*., 2017), and the B, C and D haplotypes containing additional allelic variants designated as *b3, b4* and *b5* respectively (Figure 7, Supplementary Figure S8). As for the *a* locus, we assessed variation in these genes by SNP calling against the *Pt*76 (genotype *b2*/*b3*) and 20QLD87 (*b4*/*b5*) diploid reference genomes, as well as through a *k*-mer containment analysis. This confirmed that isolates sharing the A, B, C or D haplotypes do indeed contain the same *b* locus alleles (*b2* to *b5*) as these reference haplotypes (Supplementary Table S3), in some cases along with an additional undefined divergent allele.

**Figure 7:**
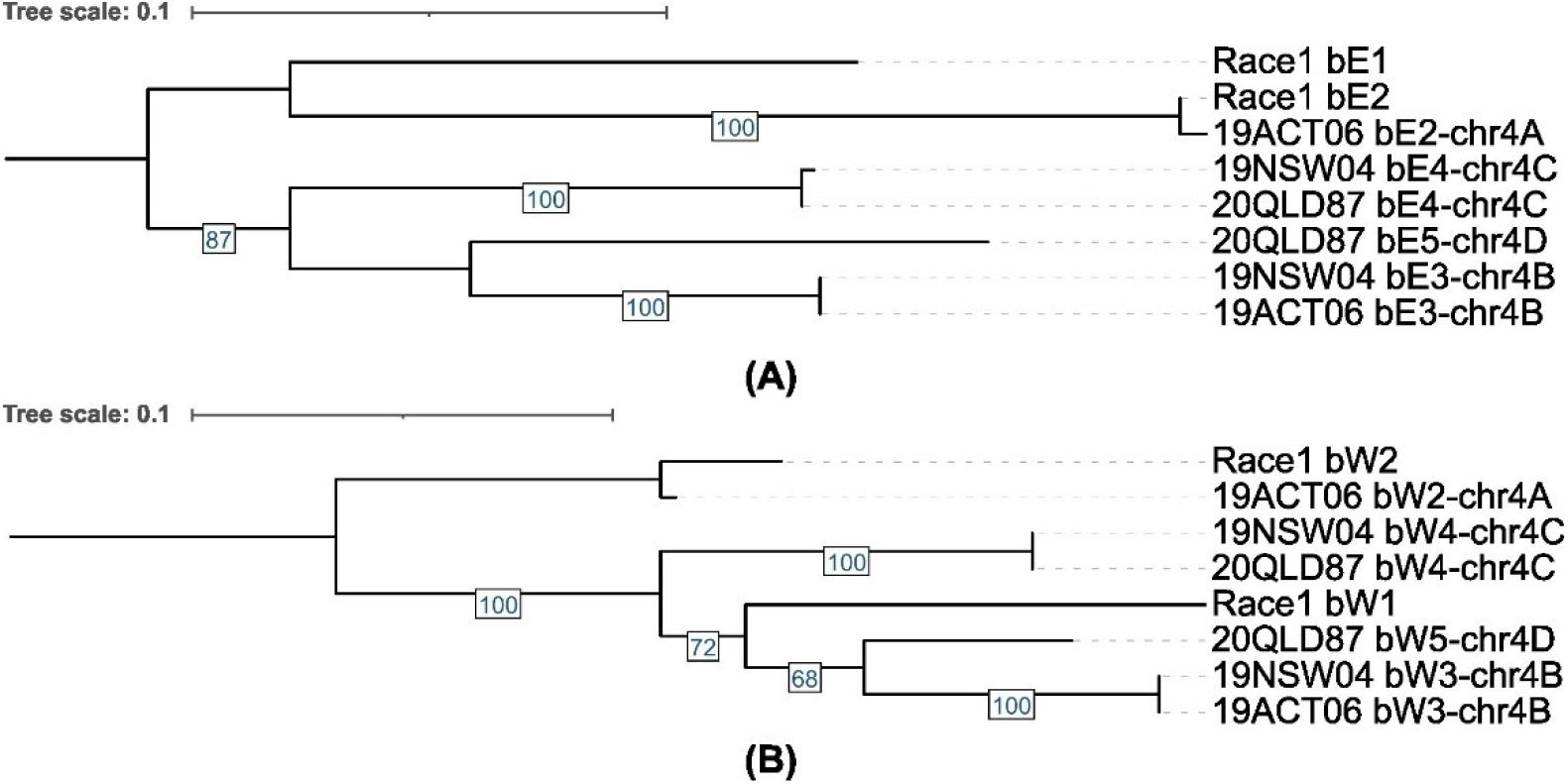
Phylogenetic trees of the mating type proteins bE and bW. Five variants of the *b* proteins are present in the listed *Pt* isolates. The b2 proteins in race 1 are shared with the *Pt*76 (19ACT96) haplotype A. Bootstrap values are shown.

## Discussion

Although somatic genetic exchange between rust strains was well established in laboratory infections, its contribution to population diversity in the field has been largely unknown and debates over whether such exchanges involved transfer of whole nuclei or parasexual recombination remained unresolved. Recent work showed that the devastating Ug99 lineage of wheat stem rust arose by a somatic nuclear exchange event (Li *et al*., 2019). Here we found by nuclear haplotype comparisons that extensive nuclear exchange events have occurred in the wheat leaf rust fungus *Pt* and have given rise to many of the long-term clonal lineages of this pathogen common around the world (Figure 8).

**Figure 8:**
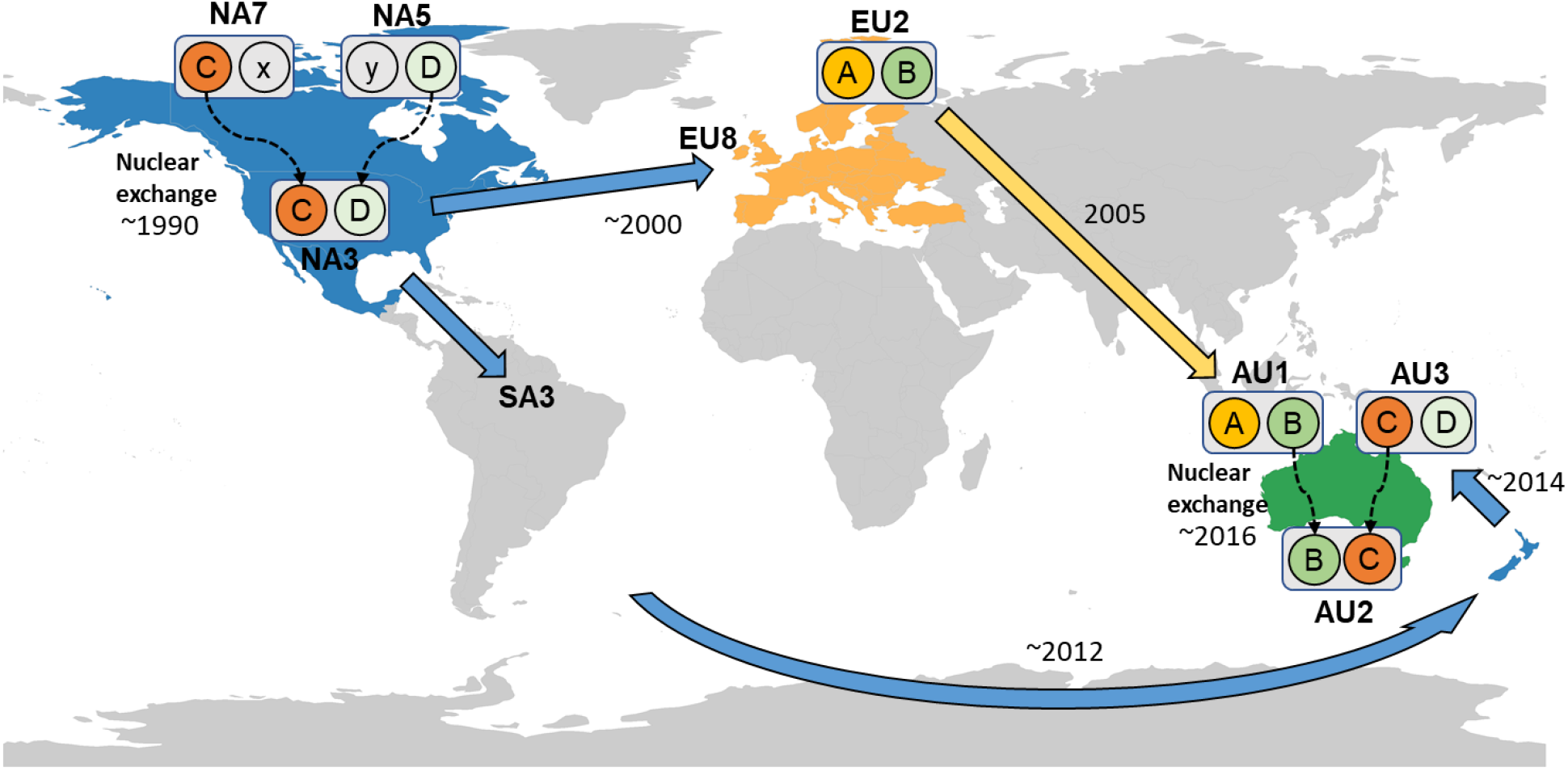
Nuclear exchange events have shaped global *Pt* lineages. The NA3 lineage most likely arose from somatic hybridisation involving an isolate of the NA5 group which donated the D genome. The NA7 clonal group contains the C haplotype and could represent the parental lineage donating this haplotype to NA3. The NA3 lineage subsequently spread to other parts of the world, including Australia. The AU2 lineage (BC) likely arose from somatic hybridization of the AB lineage (EU2) with the CD (AU3) lineage.

Detection of nucleus exchange events requires generation of nuclear-resolved genome assemblies using chromatin contact information from Hi-C sequencing to detect phase switches in assembled contigs and assign contigs to their nucleus of origin (Li *et al*., 2019). Long read assembly algorithms generate contigs that can include phase switches that require correction post-assembly (Nurk *et al*., 2020; Cheng *et al*., 2021; Rhie *et al*., 2021). For example, the PacBio Canu assemblies of *Pgt*21-0 and *Pca*203 contained 31 and 33 phase switch contigs, respectively (Li *et al*., 2019; Henningsen *et al*., 2022), while the PacBio-HiFi HiCanu assembly of *Pt*76 had 17 phase switch contigs and a hifiasm assembly of the same data contained 14 phase-switch contigs (Duan *et al*., 2022). However, the latest version of hifiasm can incorporate Hi-C data into the assembly process (Cheng *et al*., 2021) and we found that this resulted in only 0 to 3 phase switches in the three leaf rust isolate genome assemblies. Unexpectedly, the hifiasm haplotype assignments also almost perfectly matched the nuclear origin of the sequences with only 0 to 2 incorrect contig assignments per assembly. This high phasing and nuclear assignment accuracy greatly facilitates the rapid assembly of nuclear haplotypes, with the few errors readily detected and corrected using NuclearPhaser (Duan *et al*., 2022). Another assembly pipeline that can incorporate Hi-C data is FALCON-Phase (Kronenberg *et al*., 2021), which was used to generate a chromosome-level and haplotype-separated assembly for *Pt*64 (Wu *et al*., 2021). However, in this case neither contig-phasing nor nuclear origin of contigs was assessed post-assembly so the final pseudo-haplotypes may be composed of randomly assigned chromosomes from each nucleus.

Comparison of the nuclear haplotype genomes of three Australian *Pt* isolates revealed shared nuclei between these isolates, suggesting that the AU2 lineage (BC nuclear genotype) is derived from a nuclear exchange event between members of other two lineages (AB and CD nuclear haplotypes) (Figure 8). This is consistent with the known detection of these lineages in detailed pathotype surveys in Australia. In 2005, a *Pt* isolate with a novel pathotype (76-3,5,7,9,10) was first detected in Australia as a postulated exotic incursion (Huerta-Espino *et al*., 2011; Park, 2015) and several isolates derived by single-step mutations were subsequently detected (Bariana *et al*., 2022), including isolates with pathotypes matching isolates 19ACT06 and 19QLD08 (AU1 lineage, AB genotype). Similarly, an exotic incursion with pathotype 104-1,3,4,6,7,9,10,12 was first detected in New Zealand in 2012, followed by a mutational derivative with virulence to *Lr28* (pathotype 104-1,3,4,6,7,8,9,10,12) in 2014 in both New Zealand and Australia (Warren *et al*., 2018). This pathotype matches the 20QLD87 isolate (AU3 lineage, CD genotype). Subsequently, a *Pt* isolate with the novel pathotype 104-1,3,4,5,7,9,10,12 was first detected in 2016, and is now the predominant pathotype (Bariana *et al*., 2022). This pathotype matches that of 20ACT90 in the proposed hybrid AU2 lineage with the BC genotype, while the other AU2 isolates 19NSW04, 19ACT07 and 20QLD91 differ by a single virulence to Lr27/31 representing a single virulence mutation. Thus, the simplest explanation for these relationships is that the BC genotype arose by somatic hybridisation and nuclear exchange between isolates of the AB and CD genotype in Australia (Figure 8).

*Pt* is a widely distributed fungus that shows asexual reproduction in most parts of the world (Bolton *et al*., 2008) with a number of clonal groups common to Europe, North America and Africa (Kolmer, 2019). Comparison of the Australian isolates to a set of global isolates with whole genome sequence data revealed the likely origins of the AB and CD nuclear genotype lineages in Europe and North America respectively (Table 3 and Figure 4). The Australian isolate 20QLD87 (CD genotype) occurred in a clonal clade with isolates from the NA3 lineage, which has been dominant in North America since 1996 (Kolmer, 1999, 2019; Ordoñez & Kolmer, 2009; Fellers *et al*., 2021). Illumina reads from the NA3 isolates also showed full coverage of the C and D haplotype *k*-mers, confirming this relationship. The two AU1 isolates (AB genotype) form a clonal group with an isolate collected in Turkey in 2009 (09TUR23-1) and Illumina reads from this isolate fully contained all the *k*-mers of both the A and B genomes. This isolate is a member of the EU2 clade (Kolmer *et al*., 2013) and a phylogenetic analysis of shared SNP positions in a GBS data set (Kolmer, 2019) confirmed that other EU2 clade isolates also grouped with AU1 in a single clonal lineage. Although these EU2 isolates were collected in 2009, previous studies identified isolates of this pathotype group in Europe in the 1990s (Park & Felsenstein, 1998; Goyeau *et al*., 2006; Hanzalová *et al*., 2008), suggesting that it was present prior to its first detection in Australia in 2005. In addition, a set of central Asian isolates (clade CA-1) collected in 2002 and 2003 also grouped with EU2 and AU1 based on GBS data (Kolmer, 2019).

Haplotype-specific phylogenies and *k*-mer coverage analysis also provided evidence for numerous nuclear exchange events underlying diversity between major clonal lineages of *Pt* (Figures 5 and S3 to S6 and Table 4). For instance, the North American lineages NA4, NA5 and NA6 all shared the D haplotype, while NA7 contains the C haplotype and two different European isolates (FR56 and CZ10-09) contain the A haplotype, (all in combination with different alternate haplotypes). FR56 is closely related to the AU5 group of Australian isolates that also shares the A haplotype, including isolate S423 collected in 1984 and representing the founder of this clonal lineage, which dominated in Australia from 1989 to 2008 (Park *et al*., 1995; Huerta-Espino *et al*., 2011). Several of the North American clonal lineages have persisted over many decades, with NA1 including an isolate of race 1 collected in 1954, NA5 an isolate of race 5 from 1959, NA6 an isolate from 1965 (Ordoñez & Kolmer, 2009; Fellers *et al*., 2021) while previous SSR genotype analysis placed an isolate of race 9 from 1959 in NA4 (Ordoñez & Kolmer, 2009). Isolates of races 1, 5 and 9 were first described in 1920/1921 when rust pathotyping was established in the US (Johnston *et al*., 1968) suggesting these lineages were already prevalent a hundred years ago. However, isolates of the NA3 group were not detected until 1996 as a newly emerged pathotype that included the then novel combination of virulence for *LrB, Lr3bg* and *Lr17* with avirulence for *Lr28* (Kolmer, 1999), and this is now the most commonly isolated pathotype group in US surveys (Kolmer, 2019). Our data show that NA3 most likely arose from a somatic hybridisation event involving an isolate of the NA5 group which donated the D genome. The placement of the NA3 group within the NA5 clade in the phylogenetic tree derived from the D haplotype indicates that its D genome arose after the establishment of the NA5 lineage. Two other North American isolates (defined here as the NA7 clonal group) contain the C haplotype and could represent the parental lineage donating this haplotype to NA3. This is consistent with their basal position in the C haplotype phylogeny, but these two isolates were both collected in 2011 so it is also possible that they are derived from NA3. Ordoñez *et al*. (2010) found that the NA3, NA4 and NA5 lineages also occurred in South America, with the first NA3 isolates detected in 1999 in Uruguay, while NA4 and NA5 isolates dated back to at least 1981 and these shared lineages between North and South America were later confirmed by Kolmer *et al*. (2020) using GBS data. (Ordoñez *et al*., 2010) proposed that the NA3 group may have migrated to the Northern US and South America from Mexico, as similar pathotypes had been detected there earlier in the 1990s. Given the prevalence of the NA5 group in both North and South America prior to this time, it remains plausible that the hybridisation event giving rise to the NA3 lineage occurred in Mexico in the 1990s. Isolates with similar pathotype to NA3 were also detected in France in 2000, likely having spread from the Americas (Kolmer *et al*., 2013), and some isolates from the EU8 group in Europe (collected in 2004 and 2009) clustered with NA3 by both SSR and GBS SNP genotypes ((Kolmer, 2019), Figure 6 and Supplementary Figure S7). Similar genotypes were also detected in samples from Pakistan in 2010-2014 (Figure 6), indicating that this hybrid lineage has spread worldwide.

McTaggart *et al*. (2022) proposed that an alternative hypothesis to explain why two dikaryotic rust isolates would share a common nucleus is through a duplicated fertilization event, where nectar from one pycnial infection of a single haploid genotype (haplotype A), was transferred to receptive hyphae of two different pycnial infections (haplotypes B and C), giving rise to dikaryotic sexual progeny of haplotypes AB and AC. However, this is an untenable explanation for the multiple shared haplotypes detected here as it would require that all these lineages were generated by simultaneous cross-fertilisation on an alternate host plant over a hundred years ago and subsequently persisted as globally dominant races. This is not consistent with the more recent emergence of the CD and BC lineages in the 1990s and 2010s compared to the NA4, NA5 and NA6 lineages dating date back to at least the 1950s, and probably prior to the 1920s. The phylogenetic data also supports different divergence times of the common haplotypes in these lineages, rather than divergence from a single common ancestor. The duplicated fertilization hypothesis is also inadequate to explain the shared haplotypes observed between *Pgt*21, Ug99 and at least three other lineages in wheat stem rust for similar reasons (Li *et al*., 2019). On the other hand, the plethora of laboratory evidence for somatic exchange after infection of host plants by two different rust genotypes (Park & Wellings, 2012) suggests that this is not a rare or unlikely event. Therefore, the most parsimonious explanation for the relationships between the *Pt* and *Pgt* isolates with shared haplotypes is that they arose via somatic hybridisation and nuclear exchange events.

Early studies on laboratory induced somatic exchange were hampered by the lack of molecular markers, which prevented accurate discrimination between simple exchange of nuclei of opposite mating-type, versus more extensive parasexual recombination. In some cases, only two non-parental phenotypic classes were observed (Flor, 1964; Bartos *et al*., 1969), consistent with the former model. Flor (1964) conducted the most genetically controlled analysis using isolates with known nuclear genotypes for several Avr loci and showed that all the non-parental isolates recovered had no recombination between these genetic markers, suggesting in the inheritance of an intact nucleus from one somatic parent. In other cases, the isolation of additional recombinant classes suggested recombination, although this was also ascribed to isolate contamination (Barr *et al*., 1964). The haplotype resolved genome data here shows clearly that no recombination occurred in the generation of the BC genotype (AU2 lineage) in either of the parental isolates prior to donation of their nuclei, nor in the hybrid line subsequent to the exchange event. This was also the case for hybrid Ug99 strain in *Pgt* (Li *et al*., 2019). Likewise, the C and D haplotypes of NA3 derived from NA5 and NA7 remain in separate nuclei indicating no recombination in the hybrid after fusion, while the presence of the entire D haplotype in NA4 and NA6 also indicates that it was not generated by recombination in NA5 prior to the hybridisation. Similar considerations apply to the shared A haplotype in AU1 and AU5 and European isolates of *Pt*, as well as the three *Pgt* lineages sharing the B haplotype of that species. All of the *Pt* isolates related by hybridisation contain two opposite alleles (+/-) at the *a* mating type locus, consistent with this being a requirement for viable hybrid. Thus, it appears that somatic hybridisation in *Pt* and *Pgt* typically involves whole nuclear exchange without recombination.

Hybridisation and nuclear exchange events seem to be relatively common in the evolution of these dikaryotic rust species, whose populations consist of long-lived clonal lineages in most parts of the world. Indeed, it seems global populations of *Pt* and *Pgt* are dominated by isolates with different combinations of a relatively small number of haploid genotypes. This may be a consequence of the general absence of the sexual hosts in almost all wheat growing areas. Given these observations and the frequency of somatic nuclear exchange events detected here, there is the potential for repeated shuffling of haplotypes within populations. This could result in independent events giving rise to the same haplotype combinations in different locations or times and could also recreate pre-existing haplotype combinations. For example, Figure 6A suggests that some European isolates from 2009 are closely related to the AU2 (BC) group. Thus, it is possible that two different hybridisation events occurred to generate this haplotype combination independently in Europe and Australia. Alternatively, this hybrid lineage may have originated in Europe and then migrated to Australia coincidently after the AB and CD parental linages became established there. Furthermore, Figure 5 suggests that one branch of the NA5 group may contain a D nucleus derived from the NA3 (CD) group, which could result from a hybridisation between isolates of these groups effectively exchanging related but slightly diverged D genomes. Indeed, comparison of the NA3 lineage phylogenies derived from the C and D haplotypes (Figure 5 C and D) reveals several incongruities, which could occur if members of this lineage have undergone repeated exchanges of the C and D nuclei. Generating haplotype phased genome references for additional global isolates will help to confirm the proposed origins of nuclear haplotypes in these strains and identify the other haplotypes present and potential exchange events involving those haplotypes.

## Materials and Methods

### Sampling and pathotyping of the *Puccinia triticina* isolates

Rust infected samples from wheat cultivar Morocco were collected in 2019 and 2020 from the CSIRO field site in Canberra, ACT (19ACT07 and 20ACT90) and from the wheat cultivar Grenade in a field at the Department of Primary Industries, Wagga Wagga, NSW (19NSW04). Three samples were collected in 2019/20 from an unknown wheat cultivar in Warwick, QLD (20QLD87) or Gatton QLD (19QLD08 and 20QLD91). *Puccinia triticina* cultures were purified through single pustule isolation and pathotyped using the standard Australian wheat differential sets carrying unique resistance genes and nomenclature for leaf rust (McIntosh *et al*., 1995) (Supplementary Table S4).

### PacBio HiFi DNA and Hi-C sequencing

High molecular DNA from urediniospores was extracted as previously described (Li *et al*., 2019). DNA quality was assessed with a Nanodrop Spectrophotometer (Thermo Scientific, Wilmington, DE, USA) and the concentration quantified using a broad-range assay in Qubit 3.0 Fluorometer (Invitrogen, Carlsbad, CA, USA). DNA library preparation (10-15 Kb fragments Pippin Prep) and sequencing in PacBio Sequel II Platform (One SMRT Cell 8M) were performed by the Australian Genome Research Facility (AGRF) (St. Lucia, Queensland, Australia) following manufacturer’s guidelines. For DNA-crosslinking and subsequent Hi-C sequencing, 100 mg of urediniospores was suspended in 4 ml 1% formaldehyde and incubated at room temperature (RT) for 20 min with periodic vortexing. Glycine was added to achieve 1 g/100 ml, following a 20 min incubation period at RT with periodic vortexing. The suspension was centrifuged at 1,000 *g* for 1 min and the supernatant was removed. Spores were then washed with H2O, centrifuged at 1,000 *g* for 1 min, and the supernatant was then removed. The spores were then transferred to a liquid nitrogen-cooled mortar and ground before being stored at –80°C or on dry ice. After treatment spores were shipped to Phase Genomics (Seattle, WA, U.S.A) for Hi-C library preparation and sequencing.

### Illumina short-read whole-genome sequencing of *Pt* isolates

Genomic DNA was extracted from 30 mg of urediniospores per isolate using the Omniprep DNA isolation kit (G-Biosciences). DNA concentration was determined using Qubit 3.0 Fluorometer (LifeTechnologies) before submission for whole genome sequencing. A transposase-based library was prepared for each sample with DNA Prep (M) Tagmentation kit (Illumina) at the Australian Genome Research Facility (AGRF) (St. Lucia, Queensland, Australia) following manufacturer’s guidelines. DNA sequencing was completed at AGRF using a NovaSeq S4, 300 cycles platform (Illumina) to produce 150 bp paired-end reads.

### Genome assembly and scaffolding

The HiFi reads of the isolates 19NSW04 and 20QLD87 were assembled with hifiasm 0.16.1 in Hi-C integration mode and with default parameters (19NSW04: 15.2 Gb HiFi reads and 34.8 Gb Hi-C reads; 20QLD87: 12.3 Gb HiFi reads and 43.9 Gb Hi-C reads) (Cheng *et al*., 2021). Contaminants were identified using sequence similarity searches (BLAST 2.11.0 -db nt -evalue 1e-5 -perc_identity 75) (Altschul *et al*., 1990). HiFi reads were aligned to the assembly with minimap2 2.22 (-ax map-hifi --secondary=no) (Li, 2018) and contig coverage was called using bbmap’s pileup.sh tool on the minimap2 alignment file (http://sourceforge.net/projects/bbmap/). All contaminant contigs, contigs with less than 5x coverage and the mitochondrial contigs were removed from the assembly. BUSCO completeness was assessed with version 3.0.2 (-l basidiomycota_odb9 -sp coprinus) and Augustus parameters pre-trained on the *Pt*76 assembly (Simão *et al*., 2015). The HiFi reads of the *Pt*76 isolate (Duan *et al*., 2022) were re-assembled with hifiasm 0.16.1 in Hi-C integration mode and with default parameters (Cheng *et al*., 2021) to assess improvement in phasing compared to the previously published HiCanu assembly (Duan *et al*., 2022). Phasing of the assembled haplotypes was confirmed with the NuclearPhaser pipeline version 1.1 (https://github.com/JanaSperschneider/NuclearPhaser) (Duan *et al*., 2022). The HiFi reads of the 20QLD87 isolate were also assembled with HiCanu 2.2.0 to confirm phase switch boundaries (genomeSize=120m -pacbio-hifi) (Nurk *et al*., 2020) and contigs were aligned to the hifiasm assembly with minimap2 2.18 (Li, 2018).

We curated nuclear-phased chromosomes for each assembly by scaffolding the two haplotypes separately and then further joined scaffolds into chromosomes through visual inspection of Hi-C contact maps. For scaffolding of the individual haplotypes, the Hi-C reads were mapped to each haplotype using BWA-MEM version 0.7.17 (Li & Durbin, 2009) and alignments were then processed with the Arima Genomics pipeline (https://github.com/ArimaGenomics/mapping_pipeline/blob/master/01_mapping_arima.sh). Scaffolding was performed using SALSA version 2.2 (Ghurye *et al*., 2019). Hi-C contact maps were produced with HiC-Pro 3.1.0 (MAPQ = 10) (Servant *et al*., 2015) and Hicexplorer 3.7.2 (Ramírez *et al*., 2018).

### Gene prediction and repeat annotation

De novo repeats were predicted with RepeatModeler 2.0.2a and the option -LTRStruct (cite https://pubmed.ncbi.nlm.nih.gov/32300014/). RepeatMasker 4.1.2p1 (-s -engine ncbi) (http://www.repeatmasker.org) was run with the RepeatModeler library to obtain statistics about repetitive element content. For gene prediction, RepeatMasker was run with the RepeatModeler library and the options -s (slow search) -nolow (does not mask low_complexity DNA or simple repeats) -engine ncbi. RNAseq reads from *Pt*76 (Duan *et al*., 2022) were aligned to the genome with HISAT2 (version 2.1.0 --max-intronlen 3000 --dta) (Kim *et al*., 2019) and genome-guided Trinity (version 2.8.4 --jaccard_clip --genome_guided_bam --genome_guided_max_intron 3000) was used to assemble transcripts (Grabherr *et al*., 2011). We then aligned each RNAseq sample to the individual haplotype chromosomes as well as the unplaced contigs with HISAT2 (version 2.1.0 --max-intronlen 3000 --dta) (Kim *et al*., 2019). We used StringTie 2.1.6 (-s1 -m50 -M1) to assemble transcripts for each sample (Pertea *et al*., 2015). The transcripts of the ungerminated and germinated spore samples were merged into a spore transcript set for each haplotype chromosome as well as the unplaced contigs with StringTie (--merge). The transcripts of the infection time point samples were merged into an infection transcript set for each haplotype chromosome as well as the unplaced contigs with StringTie (--merge).

Funannotate 1.8.5 (Palmer & Stajich, 2020) was run to train PASA (funannotate update) with the preassembled Trinity transcripts as input (Haas *et al*., 2008). CodingQuarry 2.0 (Testa *et al*., 2015) was run in pathogen mode, once on the infection transcripts and once on the spore transcripts. For the infection transcripts, we merged the predicted genes, the predicted pathogen genes and the predicted dubious gene set into the final CodingQuarry infection gene predictions. For the spore transcripts, we merged the predicted genes, the predicted pathogen genes and the predicted dubious gene set into the final CodingQuarry spore gene predictions. We then ran funannotate predict (--ploidy 2 --optimize_augustus -- busco_seed_species ustilago --weights pasa:10 codingquarry:0) and supplied Trinity transcripts and *Pucciniomycotina* EST clusters downloaded from the JGI MycoCosm website (http://genome.jgi.doe.gov/pucciniomycotina/pucciniomycotina.info.html). We also supplied our CodingQuarry predictions to funannotate with the option -other_gff and set the weight of the CodingQuarry infection gene predictions to 20 and the weight of the CodingQuarry spore gene predictions to 2. Lastly, we also supplied the StringTie infection transcripts with weight 20 and the StringTie spore transcripts with weight 2. After the funannotate gene predictions, we ran funannotate update followed by an open reading frame prediction to capture un-annotated genes that encode secreted proteins. First, we ran TransDecoder 5.5.0 (https://github.com/TransDecoder/TransDecoder) on the StringTie infection transcripts (TransDecoder.LongOrfs -m 50 and TransDecoder.Predict --single_best_only). We selected ORFs that have a start and stop codon (labelled as ‘complete’) and predicted those that have a signal peptide (SignalP 4.1 -u 0.34 -U 0.34) and no transmembrane domains outside the N-terminal signal peptide region (TMHMM 2.0) (Krogh *et al*., 2001; Petersen *et al*., 2011). We added genes encoding secreted proteins to the annotation using agat_sp_fix_overlaping_genes.pl (Dainat *et al*., 2022), which creates isoforms for genes with overlapping coding sequence. In line with funannotate, we did not include genes encoding secreted proteins that are >90% contained in a repetitive region in the final annotation.

### Genome comparisons and *k*-mer containment screening

The haplotype chromosomes were compared to each other with mummer 4.0.0rc1, using nucmer and dnadiff (Marçais *et al*., 2018). Genomic dot plots were produced with D-GENIES (Cabanettes & Klopp, 2018). Mash 2.3 (Ondov *et al*., 2019) was used for *k*-mer containment screening (mash screen).

### Phylogenetic trees and mating type loci

Illumina reads were downloaded from NCBI and cleaned with trimmomatic v0.38 (Bolger *et al*., 2014) and then aligned against the diploid genome assemblies with bwa mem 0.7.17 (Li & Durbin, 2009). The alignment files of our seven isolates were filtered for minimum quality 30 as the coverage was substantially higher than for the alignments of the other global isolates. SNPs were called with FreeBayes 1.3.5 (--use-best-n-alleles 6) in parallel mode (Garrison & Marth, 2012) against the diploid chromosomes and the individual haplotypes. SNPs were filtered using vcffilter of VCFlib 1.0.1 (https://github.com/vcflib/vcflib) with the parameter -f “QUAL > 20 & QUAL / AO > 10 & SAF > 0 & SAR > 0 & RPR > 1 & RPL > 1 & AC > 0”. Bi-allelic SNPs were selected with vcftools (--min-alleles 2 --max-alleles 2 --max-missing 0.9 --maf 0.05) (Danecek *et al*., 2011) and converted to multiple sequence alignment in PHYLIP format using the vcf2phylip script (Ortiz, 2019). Phylogenetic trees were constructed using RAxML v. 8.2.12 (Stamatakis, 2014). We generated 500 bootstrap trees (-f a -# 500 -m GTRCAT) and a maximum-likelihood tree (-D) and incorporate these models into a final tree (-f b -z -t -m GTRCAT). Phylogenetic trees were visualized in iTOL v6 (Letunic & Bork, 2007) with the isolate ISR850 as the outgroup. We used the publicly available GBS SNP vcf file (https://conservancy.umn.edu/handle/11299/208672) and the genomic Illumina data to build a phylogenetic tree as follows. First, we mapped the clean Illumina reads to the *Pt* ASM15152v1 draft assembly downloaded from https://fungi.ensembl.org/Puccinia_triticina/Info/Index (Kolmer *et al*., 2020). SNPs were called as described in the previous paragraph. We intersected the GBS SNP vcf file with the Illumina SNP file using bcftools isec (Danecek *et al*., 2021) and kept SNPs shared by both files. Phylogenetic trees were constructed as described in the previous paragraph.

The b1-b5 proteins were aligned with mafft 7.4.90 (Katoh & Standley, 2013) and a phylogenetic tree was built with iqtree2 2.2.0.8 (-B 1000 -alrt 1000) (Minh *et al*., 2020) and visualized and midpoint-rooted in iTOL v6 (Letunic & Bork, 2007). SNP statistics were collected with bcftools stats and coverage statistics with samtools coverage (Danecek *et al*., 2021).

## Data availability

All sequence data and assemblies generated in this study are available at the NCBI BioProject PRJNA902835 (https://www.ncbi.nlm.nih.gov/bioproject/PRJNA902835). Sequencing reads, assemblies and gene annotation files are also available at the CSIRO Data Access Portal (https://data.csiro.au/collection/csiro:57097).

## Authors’ contributions

JS analysed and interpreted all data sets and wrote the manuscript. TH processed sequencing data and ran phylogenetic tree analysis. DCL performed pathotyping and prepared DNA samples for sequencing. SP performed pathotyping. AWM and LTH provided leaf rust samples. RM built rust collections and performed pathotyping. PND conceived the study, analysed and interpreted all data sets and wrote the manuscript. MF built rust collections, conceived the study, interpreted all data sets and wrote the manuscript. All authors contributed to manuscript writing and read and approved the final manuscript.

## Acknowledgements

We thank Dr. Narayana Upadhyaya for assistance in downloading of sequencing reads.

