## Supplementary Material for "Extensive somatic nuclear exchanges shape global populations of the wheat leaf rust pathogen *Puccinia triticina*"

**Supplementary Table S1: Statistics for the hifiasm-haplotype 1 and hifiasm-haplotype 2 assemblies before scaffolding.**

| Statistic | 19NSW04 hifiasm-haplotype 1 | 19NSW04 hifiasm-haplotype 2 | 20QLD87 hifiasm-haplotype 1 | 20QLD87 hifiasm-haplotype 2 |
|---:|---:|---:|---:|---:|
| Assembly size | 129.487 Mb | 123.558 Mb | 123.181 Mb | 125.281 Mb |
| # of contigs | 119 | 61 | 81 | 64 |
| N/L50 | 8/6.813 Mb | 9/6.623 Mb | 9/5.992 Mb | 9/5.943 Mb |
| Maximum scaffold length | 9.616 Mb | 8.623 Mb | 8.313 Mb | 8.988 Mb |
| GC content | 46.54% | 46.57% | 46.59% | 46.62% |
| Complete BUSCOs (%) | 96.3% | 95.2% | 95.7% | 96.3% |
| Duplicated BUSCOs (%) | 5.4% | 3.9% | 3.6% | 4.2% |
| Fragmented BUSCOs (%) | 2.7% | 2.6% | 3% | 2.7% |

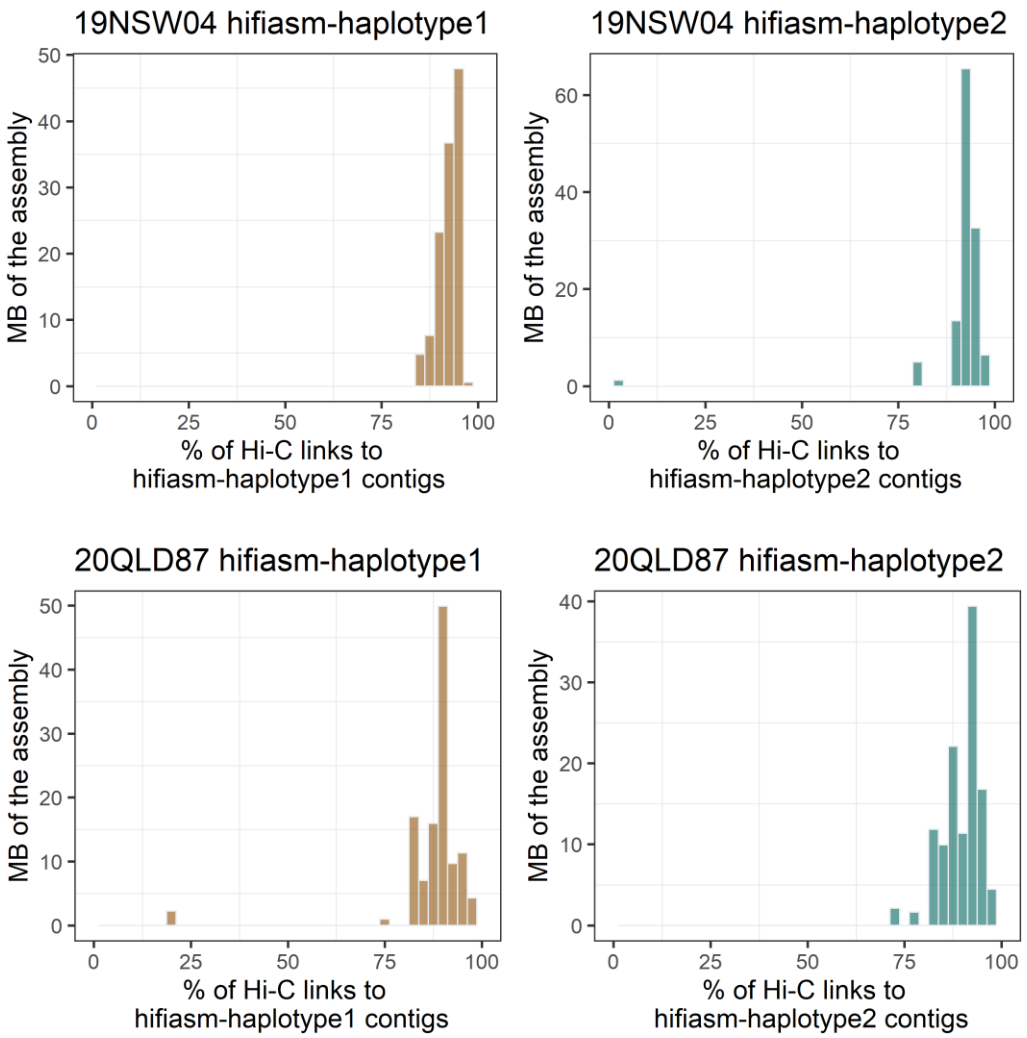


**Supplementary Figure S1: Hi-C *trans* contacts between the hifiasm-haplotype1 and hifiasm-haplotype2 in the 19NSW04 and 20QLD87 assemblies.** The two assemblies exhibit a strong dikaryotic phasing signal. In both cases the haplotype1 and haplotype2 assemblies are close to perfectly nuclear-assigned, with only two contigs larger than 150 Kb (1.2 Mb in total) assigned to the incorrect phase in19NSW04 and only a single mis-assigned contig (2.2 Mb) in 20QLD87.

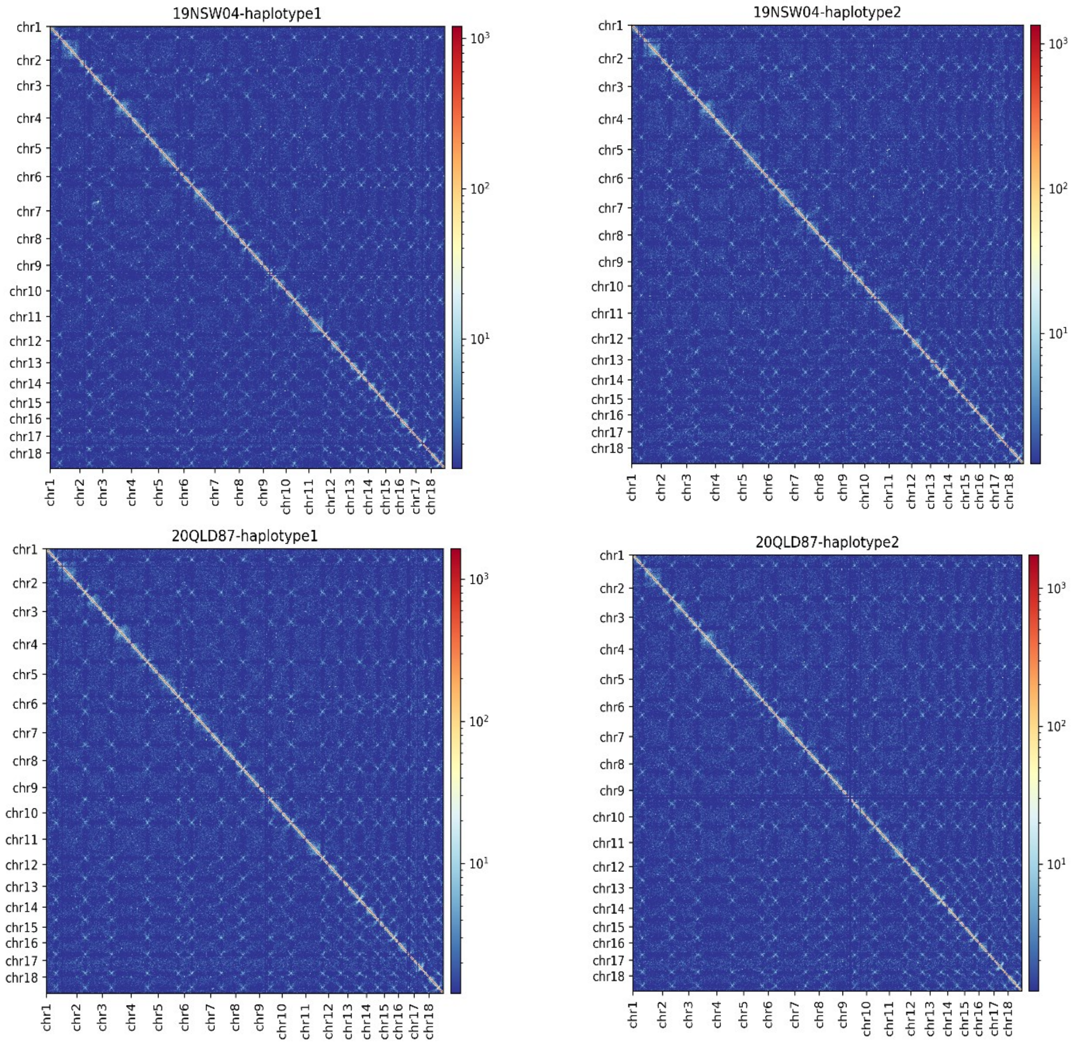


**Supplementary Figure S2: Hi-C contact maps (100 Kb resolution) for the haplotype chromosomes of the two isolates 19NSW04 and 20QLD87.** The 18 centromeres are visible as distinct outwards-spreading bowtie-like shapes.

**Supplementary Table S2: Statistics for genomic alignments between the *Pt76* and the 19NSW04/20QLD87 haplotype 1 and haplotype 2 chromosomes.**

| ***Within-isolate alignments*** | **19NSW04 haplotype 1 versus haplotype 2** | **20QLD87 haplotype 1 versus haplotype 2** | ***Pt76* haplotype A versus haplotype B** |
|---|---|---|---|
| Aligned bases | 97.7% | 97.4% | 97.1% |
| Average identity of 1-to-1 alignments | 99.5% | 99.5% | 99.5% |
| Average identity of M-to-M alignments | 99% | 99.1% | 99% |
| Translocations | 878 | 848 | 1,053 |
| Inversions | 143 | 132 | 164 |
| Insertions | 9,073 | 8,617 | 10,898 |
| Total SNPs | 329,851 | 301,814 | 334,189 |
| Total Indels | 184,447 | 167,011 | 186,643 |

| ***Between-isolate alignments*** | **19NSW04 haplotype 1 versus *Pt76* haplotype B** | **19NSW04 haplotype 2 versus *Pt76* haplotype B** | **20QLD87 haplotype 2 versus 19NSW04 haplotype 2** | **20QLD87 haplotype 2 versus 19NSW04 haplotype 1** | **20QLD87 haplotype 1 versus *Pt76* haplotype A** |
|---|---|---|---|---|---|
| Aligned bases | 97.25% | **99.21%** | 99.51% | **99.7%** | 97.4% |
| Average identity of 1-to-1 alignments | 99.49% | **99.99%** | 99.66% | **99.99%** | 99.49% |
| Average identity of M-to-M alignments | 99.06% | **99.96%** | 99.43% | **99.97%** | 99.07% |
| Translocations | 903 | **1** | 458 | **4** | 970 |
| Inversions | 150 | **6** | 90 | **2** | 148 |
| Insertions | 10,086 | **1,376** | 5,141 | **396** | 9,961 |
| Total SNPs | 326,202 | **2,966** | 219,335 | **2,226** | 327,782 |
| Total Indels | 182,284 | **9,457** | 125,084 | **6,522** | 7,948 |
| **Assignment** | | 19NSW04 haplotype 2 = B | | 19NSW04 haplotype 1 = C<br>20QLD87 haplotype 2 = C | |

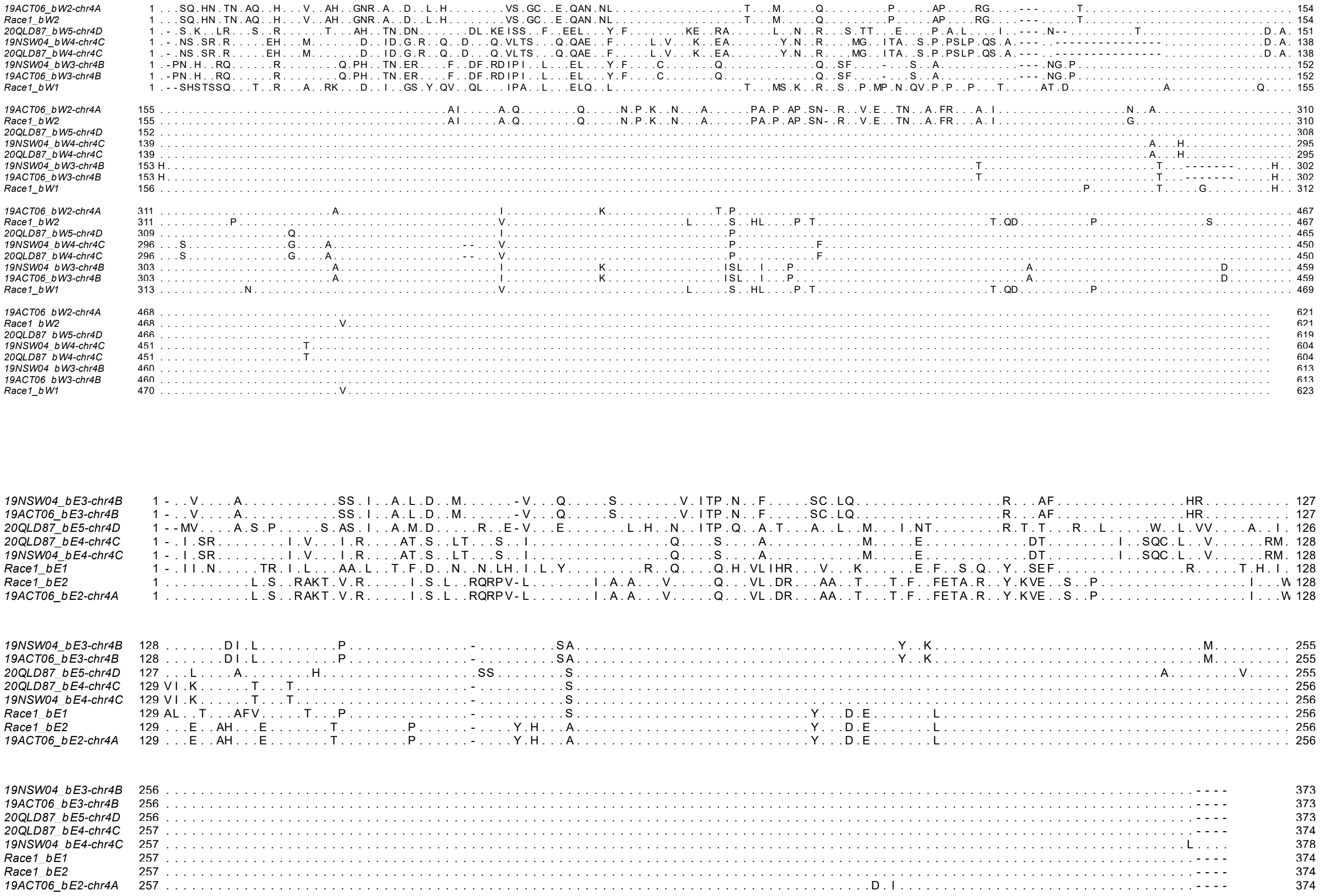

**Supplementary Figure S8: Multiple sequence alignments of the bE and bW proteins.**

| | *Pt76* b2 (haplotype A) | | | *Pt76* b3 (haplotype B) | | | 20QLD87 b4 (haplotype C) | | | 20QLD87 b5 (haplotype D) | | | |
|---|---|---|---|---|---|---|---|---|---|---|---|---|---|
| | Coverage | # hom SNPs | k-mer score | Coverage | # hom SNPs | k-mer score | Coverage | # hom SNPs | k-mer score | Coverage | # hom SNPs | k-mer score | genotype |
| AU1 (AB) | **100%** | **0** | **100%** | **100%** | **0** | **100%** | 70.8% | 9 | 98.3% | 83.4% | 24 | 98.8% | b2/b3 |
| AU2 (BC) | 62.5% | 26 | 97.9% | **100%** | **0** | **100%** | **100%** | **0** | **100%** | 82.3% | 34 | 98.8% | b3/b4 |
| 20QLD87 (CD) | 62.4% | 8 | 97.9% | 79.6% | 16 | 98.3% | **100%** | **0** | **100%** | **100%** | **0** | **100%** | b4/b5 |
| AU5 (A) | **100%** | **0** | **100%** | 67.5% | 31 | 98.1% | 66.8% | 16 | 98.4% | 75.3% | 21 | 99.1% | b2/? |
| AU4 | 60.7% | 8 | 98.2% | 72% | 15 | 98.5% | 60.2% | 16 | 98.5% | **100%** | **0** | **100%** | b5/? |
| 09TUR23-1 (AB) | **100%** | **0** | **100%** | **100%** | **0** | **100%** | 63% | 6 | 99% | 99.8% | 8 | 99.8% | b2/b3 |
| CZ10-9 (A) | **100%** | **0** | **100%** | 74.9% | 21 | 98.8% | 64.8% | 8 | 99% | 94.8% | 15 | 99.6% | b2/? |
| FR56 (A) | **100%** | **0** | **100%** | 87.4% | 19 | 99.2% | 69.6% | 12 | 98.9% | 76.6% | 15 | 99.3% | b2/? |
| EU1 | 74.6% | 10 | 97.9% | 74.3% | 15 | 97.9% | 56.7% | 11 | 98.3% | 96.4% | 7 | 99.2% | ?/? |
| EU4 | **100%** | **0** | **100%** | **100%** | **0** | **100%** | 62.9% | 9 | 98.3% | 75.5% | 22 | 98.8% | b2/b3 |
| EU5 | 91.8% | 5 | 99.5% | 74.7% | 21 | 98.7% | 76.2% | 9 | 99.1% | 94.6% | 13 | 99.6% | ?/? |
| NA1 | 98.2% | 9 | 99.2% | 71.1% | 22 | 97.7% | 64.9% | 8 | 98.2% | 75% | 23 | 98.3% | ?/? |
| NA2 | 89.2% | 8 | 99.1% | 72.2% | 22 | 98% | 67.8% | 11 | 98.5% | 81.3% | 17 | 98.8% | ?/? |
| NA3 (CD) | 63.8% | 8 | 98.1% | 69.4% | 16 | 98.3% | **99.9%** | **0** | **100%** | **99.9%** | **0** | **100%** | b4/b5 |
| NA4 (D) | 66.6% | 7 | 98.7% | 66.7% | 18 | 98% | 51.1% | 6 | 98.6% | **100%** | **0** | **100%** | b5/? |
| NA5 (D) | 70.6% | 9 | 98.5% | 74.1% | 18 | 98.7% | 69.5% | 14 | 99% | **100%** | **0** | **100%** | b5/? |
| NA6 (D) | 65.9% | 11 | 98.2% | 74.3% | 20 | 98.6% | 61.3% | 19 | 98.7% | **100%** | **0** | **100%** | b5/? |
| NA7 (C) | 72.2% | 8 | 99% | 64.6% | 14 | 98.3% | 99.9% | **0** | 99.9% | 79.9% | 8 | 99.2% | b4/? |
| Durum | 76.7% | 14 | 98.2% | 70.6% | 16 | 98% | 73.3% | 10 | 98.4% | 75.4% | 20 | 98.5% | ?/? |
| Middle East | **100%** | **0** | **100%** | 64.1% | 20 | 98.3% | **100%** | **0** | **100%** | 61.4% | 17 | 98.8% | b2/b4 |

**Supplementary Table S3:** Illumina read coverage, number of homozygous SNPs and *k*-mer containment score for the genomic loci of the b2, b3, b4 and b5 genes.

| | Differential number | Resistance gene | Cultivar | 19NSW04 | 19ACT06 | 19ACT07 | 19QLD08 | 20QLD87 | 20ACT90 | 20QLD91 |
|---|---|---|---|---|---|---|---|---|---|---|
| International series | | Lr1 | Tarsa | V | A | V | A | V | V | V |
| | | Lr2a | Webster | A | A | A | A | A | A | A |
| | | Lr3a | Democrat | V | V | V | V | V | V | V |
| Australian series | 1 | Lr20 | Thew | V | A | V | V | V | V | V |
| | 2 | Lr23 | Gaza | A | A | A | A | A | A | A |
| | 3 | Lr14a | Spica | V | V | V | V | V | V | V |
| | 4 | Lr15 | K1483 | V | A | V | A | V | V | V |
| | 5 | Lr3ka | Klein Titan | V | V | V | V | A | V | V |
| | 6 | Lr27+Lr31 | Gatcher | V | V | V | A | V | A | V |
| | 7 | Lr17a | Songlen | V | V | V | V | V | V | V |
| | 8 | Lr28 | CS 2A/2M | A | A | A | A | V | A | A |
| | 9 | Lr26 | Mildress | V | V | V | V | V | V | V |
| | 10 | Lr13 | Egret | V | V | V | V | V | V | V |
| | 11 | Lr16 | Exchange | A | A | A | A | A | A | A |
| | 12 | Lr17b | Harrier | V | V | V | V | V | V | V |
| | 13 | Lr24 | Agent | A | V | A | V | A | A | A |

**Supplementary Table S4: Pathotyping results for the set of seven Australian isolates.**
