## Supplementary figures and images for "Extensive somatic nuclear exchanges shape global populations of the wheat leaf rust pathogen *Puccinia triticina*"

### Supplementary Figure S3

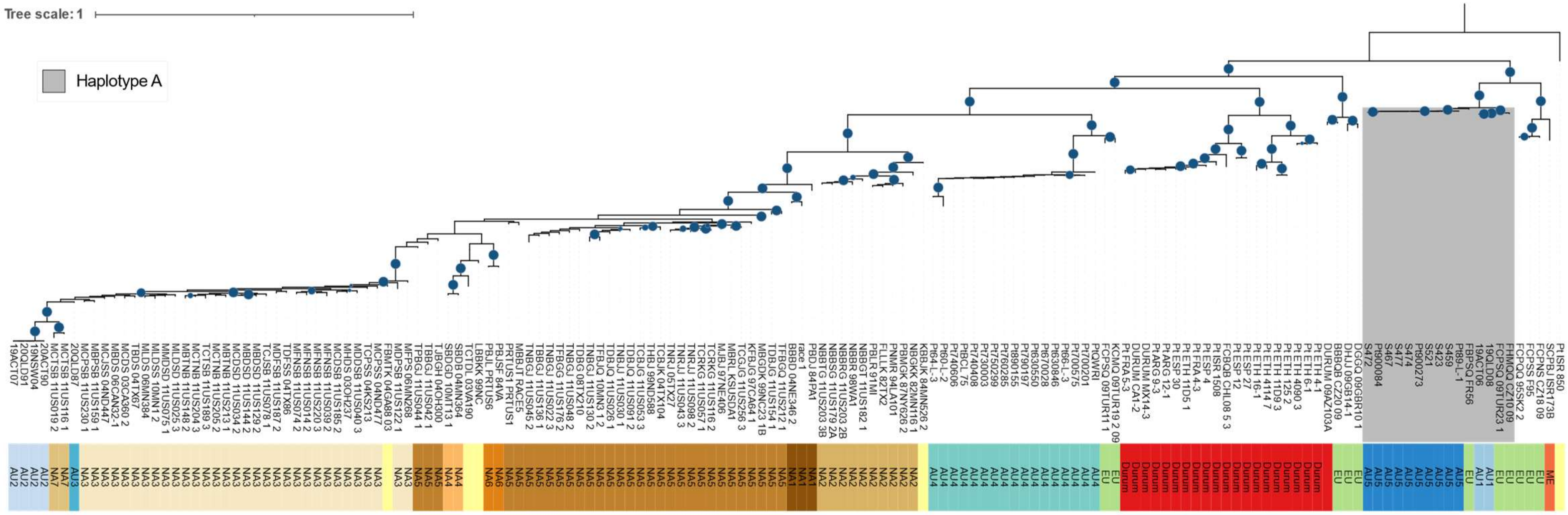


**Supplementary Figure S3: Phylogenetic tree of global isolates against *Pt76* haplotype A.**

### Supplementary Figure S4

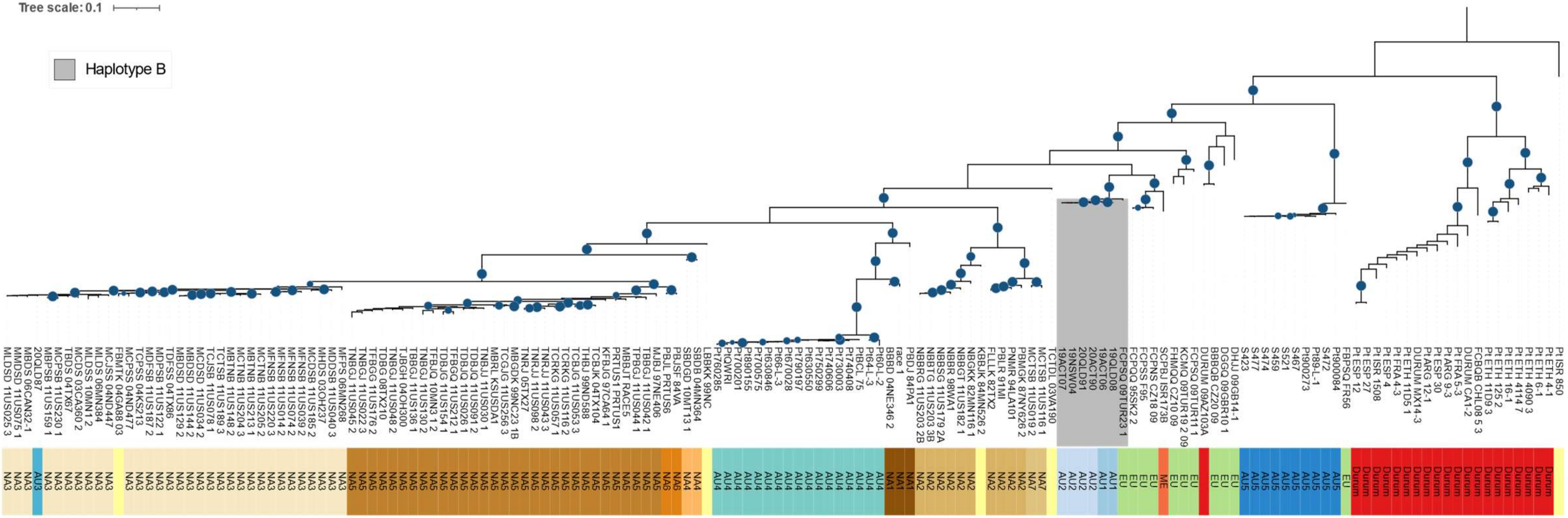


**Supplementary Figure S4: Phylogenetic tree of global isolates against 19NSW04 haplotype B.**

### Supplementary Figure S5

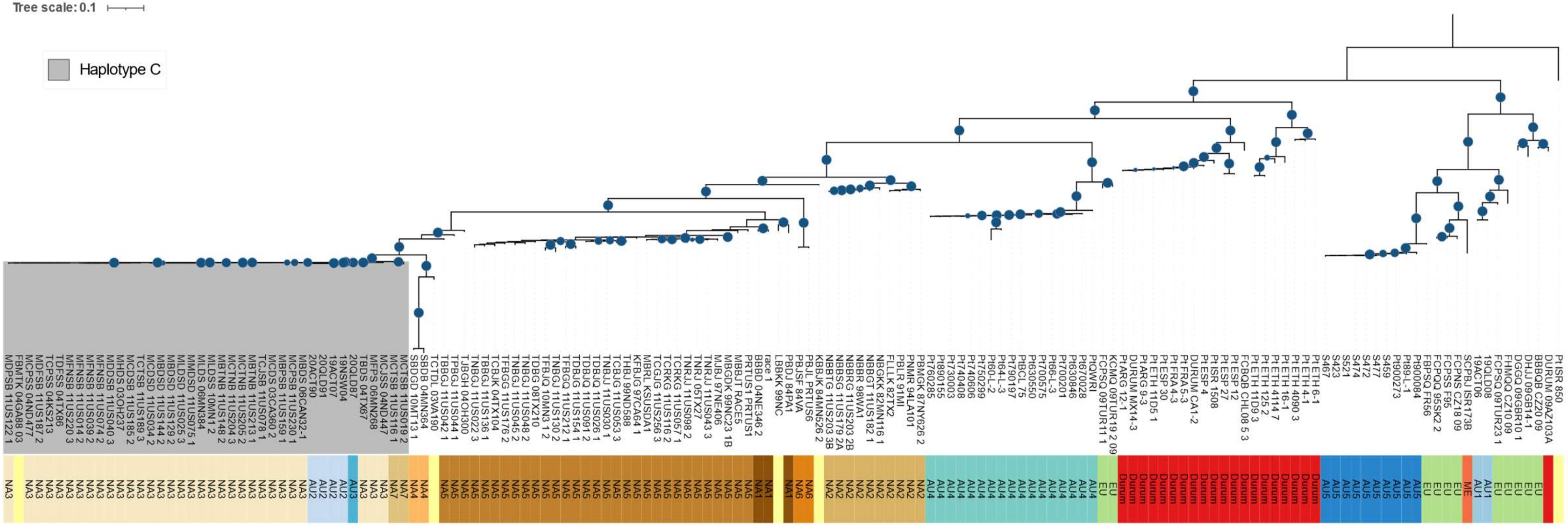


**Supplementary Figure S5: Phylogenetic tree of global isolates against 20QLD87 haplotype C.**

### Supplementary Figure S6

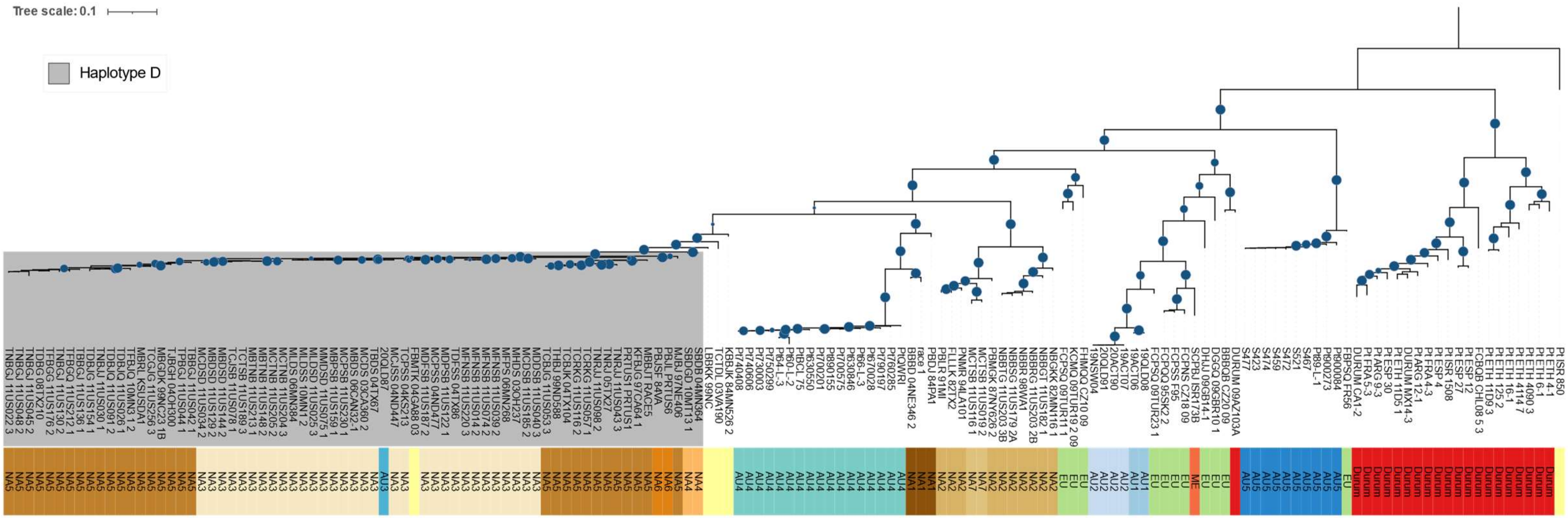


**Supplementary Figure S6: Phylogenetic tree of global isolates against 20QLD87 haplotype D.**
